# Exploring the Repertoire of Rhomboid Proteases in *Cryptosporidium parvum* Parasite: Phylogenesis, Structural motifs and Cellular Localization in Sporozoite Cells

**DOI:** 10.1101/2025.10.24.684163

**Authors:** Ilaria Vanni, Elisabetta Pizzi, Lucia Bertuccini, Alessandra Ludovisi, Fabio Tosini

## Abstract

*Cryptosporidium parvum* is an apicomplexan parasite and an important pathogen of mammals and humans, which can be infected by zoonotic transmission or directly by human-to-human contacts. This parasite attacks the small intestine, and the main symptom is a watery diarrhoea that can be particularly severe in newborns and deadly in immunodeficient subjects. Rhomboids are ubiquitous proteases embedded in cell membranes that act by cleaving other membrane proteins in or near their transmembrane domains. Apicomplexan rhomboids play an important role in approaching and invading the host cell. This study analysed the phylogenetic origin, the structural motifs and the subcellular localization of *C. parvum* rhomboids. Altogether, *C. parvum* possesses three rhomboids, namely CpRom1, CpRom2 and CpRom3. The similarity search in *Cryptosporidium* genus revealed that *C. parvum* as well as other “intestinal” species lacks a PARL-like rhomboid whereas this type of mitochondrial rhomboid was present in “gastric” species like *Cryptosporidium muris* and *Cryptosporidium andersoni.* At the genome level this was revealed by a precise excision of the PARL-like gene in intestinal species whereas the rest of chromosomal synteny was well conserved among the *Cryptosporidium* species. The analysis of the structural domains revealed that *C. parvum* rhomboids can be classified as mixed secretases and the comparison with orthologs from *Toxoplasma gondii* and *Plasmodium falciparum* showed that *C. parvum* rhomboids can be distinguished in two separate clusters based on similarities at the level of the catalytic sites. The three rhomboids were expressed simultaneously in the invasive stage of sporozoite, but each of them had a different spatial distribution. Indeed, CpRom1 had a dual localization: this rhomboid was internal at the apical complex, and it was also accumulated at the posterior pole of the sporozoite. Otherwise, CpRom2 was prevalently contained in the apical complex, and a point of accumulation was on the surface of the apical end. Differently from CpRom1 and CpRom2, CpRom3 is distributed along the entire surface of sporozoites. Finally, we listed 10 membrane proteins as candidate substrates for the *C. parvum* rhomboids based on the similarities with some proven substrates of apicomplexan rhomboids and the copresence in subcellular structures with the three rhomboids.

## Introduction

The genus *Cryptosporidium*, which belongs to the phylum Apicomplexa, is composed of many species of obligate parasites that in turn affect many species of vertebrates. *Cryptosporidium parvum* is the prevalent cause of cryptosporidiosis, a gastroenteric disease characterized by acute diarrhoea, among many mammalian species, particularly ruminants and humans (Ryan et al., 2021). Indeed, *C. parvum* is often associated to human outbreaks caused by zoonotic transmission from livestock and wild ruminants (Helmy and Hafez, 2022). It is also considered a dangerous pathogen for immunodeficient subjects, in which the infection results in a watery diarrhoea that can last for weeks, leading to a severe and sometimes deadly dehydration (Balendran et al., 2024). Moreover, *C. parvum* and the strictly related *Cryptosporidium hominis* are leading causes of mild and severe diarrhoeal cases among neonates and very young children in developing countries, accounting for thousands of deaths every year (Khalil et al., 2018). Despite its relevance for animal and human health, there are no effective treatments for this pathogen, and a viable vaccine is still to be conceived.

Rhomboids are serine-proteases provided with various transmembrane (TM) domains; a bundle composed of 6 to 7 TM helices spanning the lipidic bilayer. The rhomboid family has numerous members that are present in all kingdoms of life, from bacteria to mammals.

As proteases, rhomboids cleave other transmembrane proteins internally or near their transmembrane domain, producing two separate segments of the cleaved protein (Strisovsky, 2013). A mechanistic model of action of rhomboids is based on a three-dimensional reconstruction that shows that the TMs, those that compose the entire rhomboid domain, are organized like a sort of barrel embedded in the lipidic bilayer, with a slot on a side to allow the entry of the transmembrane domains of the target protein in the middle of the proteolytic domain, where the cleavage occurs. This process will thus produce an internal segment that will remain toward the inner side of the membrane, whereas the other protein portion will be released from the outer membrane (Strisovsky, 2013). These proteases bear a characteristic dyad as catalytic site consisting of a short consensus at TM4, namely GXSX, where X can represent various amino acids (aa), and an essential histidine placed at TM6. These amino acidic positions are conserved in all rhomboids with proteolytic activity and allow to distinguish between rhomboids and phylogenetically related but inactive pseudo-rhomboids (Tichà et al., 2018). However, other conserved amino acids flank these critical residues, so that, together with the number and the disposition of the TM domains, rhomboids can be further distinguished in different subclasses, such as secretase type A, type B and mixed secretases (Lastun et al., 2016). A further subclass is represented by PARL that are specifically linked to the inner mitochondrial membrane and that can be distinguished at the structural level for the presence of an additional TM domain at the N-terminus, thus implying that the catalytic dyad is moved from TM domains 4 and 6 to TM5 and TM7 (Lysyk et al., 2020). Obviously, rhomboids require to be allocated in a lipid bilayer to exert their proteolytic action and the cleavage mediated by rhomboids is an irreversible step in important biological processes. Thus, rhomboids mediate the EGFR signalling in *Drosophila* through the proteolysis of Spitz; the quorum sensing in bacteria like *Providencia stuartii*; the immune evasion in *Entamoeba hystolitica*; the homeostasis of mitochondrial membrane in man and other organisms (Urban and Dickey, 2011). However, thanks to the sequencing of myriads of genomes, we now know thousands of rhomboid homologs, although the function of most of them remains unknown.

The rhomboid superfamily also includes pseudoproteases, which conserve the overall structure of rhomboids but lack the enzymatic proteolytic activity. Among these we find the iRhoms, which are considered regulators of stability and trafficking of other membrane proteins as observed for human iRom2 (Dulloo et al., 2019), and the so-called derlins, which are specialized types of pseudoproteases that reside in the endoplasmic reticulum (ER) and are involved in the (ER)-associated degradation (ERAD) pathway that contributes to the degradation of defective proteins (Lemberg and Strisovsky, 2021). Derlins have also been identified in the apicoplast, which is an exclusive organelle of Apicomplexa, in *Plasmodium falciparum* and in other apicomplexans (Gallagher et al., 2011).

In general terms, apicomplexan parasites like *Toxoplasma gondii* and *P. falciparum* have a greater number of functional rhomboids if compared to mammals: 7 rhomboids in *P. falciparum* and 6 in *T. gondii* respect to the 5 identified in humans (Bergbold and Lemberg, 2013). Indeed, rhomboids in Apicomplexa play crucial roles during the invasion of the host cell. Thus, as demonstrated in *T. gondii* (Carruthers, 2006) as well as in *P. falciparum*, some rhomboids act by progressively cutting away adhesins (i.e. adhesive proteins distributed on the parasite surface) from the parasite external membrane to allow the gliding in the extracellular matrix and to favour the entry in the host cell (Sibley, 2013).

Some years ago, looking for specific antigens of *C. parvum* sporozoites, we identified a novel rhomboid in consequence of the immunogenicity of a short peptide at its N-terminus, which we referred to as CpRom; this rhomboid, composed of 990 amino acids, is the largest rhomboid ever described (Trasarti et al., 2007). More recently, investigating the functional role of this protein, we have fortuitously found out that this rhomboid, now named CpRom1, is associated with the membrane of large extracellular vesicles released by the excysted sporozoites (Bertuccini et al., 2024). A quick search by similarity with rhomboids reveals that *C. parvum* genome encodes for three functional rhomboids in total, but a comprehensive analysis of these rhomboids is still lacking. The aim of this study is the comparison of these rhomboids with the other apicomplexan rhomboids in terms of phylogenetic origin and similarity as well as of their expression in the invasive stage of sporozoites and subcellular localization. Furthermore, the high similarities of some *C. parvum* proteins with apicomplexan adhesins allowed the identification of plausible cleavage substrates for the three *C. parvum* rhomboids.

## Material and methods

### General microbiological and DNA techniques

The recombinant and microbiological techniques, media, preparation of plasmid DNA and isolation of restricted fragments all followed standard procedures (Sambrook et al., 1989). DNA sequencing was performed using Sanger’s procedure (Sanger et al., 1977).

### Sporozoite purification

Fresh C. parvum oocysts (Iowa strain) were supplied by Bunch Grass Farm (Deary, Idaho USA), stored at 4°C in PBS with penicillin (1000 U.I./ml) and streptomycin (1 mg/ml). The excystation procedure was performed as described (Bertuccini et al., 2024). Briefly, aliquots of 1×10^5^ per ml for microscopy or 1×10^7^ oocysts per ml for total lysate were pelleted at 376 × g, for 5 min, resuspended in 1 ml of 10 mM HCl, and then incubated at 37°C for 10 min. Oocysts were pelleted again as above and resuspended in 1 ml of excystation medium D-MEM (Sigma-Aldrich) plus 2 mM sodium taurocholate (Sigma-Aldrich); the mixture was maintained at 15°C, for 10 min, and then moved to 37°C to induce the excystation. Excystation mixtures were sampled at various times after the induction, and excystation stopped in ice.

### RNA extraction and RT-PCR analysis

Total RNA was extracted from 1 x 10^7^ excysted oocysts using the RNeasy mini purification kit (Qiagen Gmbh, Hilden, Germany), following the manufacturer’s protocol for yeast. The preparation was digested with Dnase RQ1 (Promega Corporation, Madison, WI, USA) for 1 h at 37 °C and purified using the RNeasy mini purification kit (Qiagen Gmbh, Hilden, Germany), following the RNA clean up protocol. The reverse transcriptase (RT) reaction was performed on 3 μg of total RNA using the primers specified in Table 1. The PCR amplification consisted of 30 cycles (60 s at 94 °C, 90 s at 55 °C and 60 s at 72 °C), followed by an extension cycle (10 min at 72 °C) on a GenAmp PCR System 2400 (Perkin-Elmer, Norwolk, CT, USA). The positive control reaction was performed, as above, on 3 μg of total RNA with primers for the *Cpa135*, which is expressed at the sporozoite stage (Tosini et al., 2004). All negative control reactions were performed on the same total RNA sample, omitting the RT reaction.

**Table 1.** List of the *C. parvum* rhomboids in this study.

| Assigned Name<br>(CryptoDB name) | CryptoDB ID | UniProt ID | Amino Acids | Chromosome localization | GO Terms | Localization in sporozoites<sup>1</sup> | OOC/SPO<sup>2</sup> | Intracell. Stages<sup>2</sup> | Proteomics | Alternative name with reference |
| --- | --- | --- | --- | --- | --- | --- | --- | --- | --- | --- |
| **CpRom1**<br>(Peptidase S54 rhomboid) | **cgd6_760** | Q5CXK3 | 990 | 6 | Proteolysis<br>Integral component of membrane<br>Serine-type endopeptidase activity | Inner membrane complex or Plasma membrane | Low | High | Oocyst Wall Proteome<sup>3</sup><br>Sporozoite Proteome<sup>4</sup> (Pept 5) | ID2/CpRom (Trasarti et al., 2007).<br>CpRom1 (Bertuccini et al., 2024) |
| **CpRom2**<br>(Rhomboid-like protein) | **cgd7_3020** | Q5CYB1 | 464 | 7 | Proteolysis<br>Integral component of membrane<br>Serine-type endopeptidase activity | unassigned<sup>5</sup> | High | Low | No Data | CpRom1 (Castellanos-Gonzalez A et al., 2019) |
| **CpRom3**<br>(Peptidase S54 rhomboid domain containing protein) | **cgd3_980** | FOX528 | 282 | 3 | Proteolysis<br>Integral component of membrane<br>Serine-type endopeptidase activity | unassigned<sup>3</sup> | Medium | High | Oocyst wall proteome<sup>6</sup> (Pept 2) | CpRom1 (Li M et al., 2016).<br><sup>7</sup>CpRom1 (Gao X et al., 2021) |

<sup>1</sup> Localization were based on results reported in Guérin A et al. (2023).
<sup>2</sup> Expression values were based on Mirhashemi ME et al. (2018)
<sup>3</sup> Proteomics data based on Truong and Ferrari (2006)
<sup>4</sup> Proteomics data based on Sanderson SJ et al. (2008)
<sup>5</sup> This protein has been identified in sporozoites, but not assigned to a specific compartment or cluster.
<sup>6</sup> Wang L et al. (2022)
<sup>7</sup> The discovery of this rhomboid is presented as novel and independent from that of Li M et al. (2016)

### Expression of recombinant rhomboid in E. coli strains and preparation of antisera

Strategy for cloning the recombinant 6His-CpRom1 has been reported in Bertuccini et al. (2024). Similarly, 6His-CpRom2 and 6His-CpRom3 were cloned using extended primers with opportune restriction sites at their ends (*Sac*I and *Sma*I for 6His-CpRom2, *Bam*HI and *Sma*I for 6His-CpRom3). The primer sequences are reported in Supplementary Table 1 and below the procedure in brief.

Both amplifications were performed with 2X Phusion Flash High fidelity PCR Master Mix (Finnzymes) using 80 ng of cDNA (see above) as template for each reaction as follows: 95°C for 5 min, 35 cycles of 94°C for 15 sec, 50°C for 15 sec, 72°C for 4 min, and a final extension at 72°C for 10 min in a Veriti 96-well thermal cycler (Applied Biosystem). Amplicons were purified with QIAquick PCR Purification Kit (Qiagen GmbH, Hilden, Germany) and digested with *Sac*I plus *Sma*I restriction enzymes (New England Biolabs) for 6His-CpRom2 and *Bam*HI and *Sma*I for 6His-CpRom2 and 6His-CpRom3, respectively. Resulting fragments were then ligated in the *Sac*I plus *Sma*I and *Bam*HI plus *Sma*I digested pQE80 vector (Qiagen GmbH, Hilden, Germany) for cloning 6His-CpRom2 and 6His-CpRom3, respectively, with the Quick Ligase kit (New England Biolabs), and the ligation mix was used to transform the Escherichia coli M15 strain. Positive clones were selected on LB agar plates with 100 µg/ml ampicillin and 25 µg/ml kanamycin by PCR screening; recombinant plasmids with recombinant CpRom2 and CpRom3 ORFs were sequenced to check the fusions with the histidine-tag coding sequences. For the purification of the two recombinant proteins, 20 ml of an overnight culture of recombinant bacteria were inoculated in 1 L of LB with 100 µg/ml ampicillin and 25 µg/ml kanamycin, and cultured at 37°C, with vigorous shacking, until a 0.6-0.8 OD was reached; then, 1 mM IPTG was added to the culture and the bacterial growth carried on for additional 3 h. Bacteria pellets were obtained by centrifugation at 1,500 × g for 10 min, then resuspended in 100 ml ice-cold PBS and centrifuged again at 3,300 × g for 20 min. The pellets were lysed in 20 ml of denaturing buffer A (100 mM NaH2PO4, 10 mM Tris-HCl, 6 M guanidine-HCl, pH 8.0), and stirred at 25°C, overnight. The lysates were clarified by centrifugation at 9,400 × g, for 30 min. Then, 5 ml of 50% Ni-NTA resin were added and gently mixed by stirring for 60 min, at 25°C; the slurries were packed in a 10 ml column and then the resin was washed with 40 ml (8 x 5 ml wash) of buffer C (100 mM NaH2PO4, 10 mM Tris-HCl, 8 M urea, pH 6.3). Purified proteins were eluted with 4 aliquots of 2.5 ml of buffer D (100 mM NaH2PO4, 10 mM Tris-HCl, 8 M urea, pH 5.9) and finally with 4 aliquots of 2.5 ml of buffer E (100 mM NaH2PO4, 10 mM Tris-HCl, 8 M urea, pH 4.5). Pooled fractions from buffer D and separately from buffer E were extensively dialyzed against PBS with increasing concentrations of glycerol (from 10% to 50%) in a Slide-A-Lizer™ (ThermoFisher) dialysis cassette with a 10 kDa cut-off of.

### Western blot, mouse sera preparation and Dot blot analysis

Solubilization of oocyst/sporozoite proteins was obtained in lysis buffer [10 mM Tris-HCl pH 7.5, DDT 10 mM, EDTA 1 mM, SDS 1%, Triton X100 1%, Sodium deoxycholate 0,5%, plus Protease Inhibitor Cocktail (Sigma Aldrich) added immediately prior the use], with the following procedure: oocyst/sporozoite pellets corresponding to 1 x 10^7^ were resuspended in 20 μl of lysis buffer, incubated at RT for 5 min in thermomixer and mixed at 1000 rpm (Eppendorf), then temperature was increased to 70° C and incubation continued for 5 min. Final denaturation was performed with the addition of 20 μl of Laemmli buffer and incubation at 95 °C for 5 min. Denatured samples were centrifuged at 12,000 rpm for 1 min before loading in SDS-PAGE. Electrophoresis was performed on MiniProtean apparatus (Biorad) with precast gels 4-15% (Biorad). Expression of recombinant proteins was checked on 3 ml of bacterial culture induced for 3 hours with 1 mM of IPTG, pelleted and lysed with Laemmli sample buffer, loaded and ran on a 4-20% TGX™ SDS-PAGE precast gels (Bio-Rad, Hercules, CA, USA) and finally tested in Western blot as below. Aliquots of 20-50 μl of purified recombinant proteins were incubated at 95 °C for 5 min in Laemmli sample buffer before loading on SDS-PAGE as above. Gels were transferred to nitrocellulose membranes (Bio-Rad), which were then blocked in TNT buffer (0.1 M Tris-Cl pH 7.5, 150 mM NaCl, 0.1% Tween-20) with 5% non-fat milk. Blots were incubated with primary antibodies for 1 h at room temperature. For recombinant 6His-CpRom2 and 6His-CpRom3, blots were incubated with mouse monoclonal RGS-His antibody (Qiagen) diluted 1:1,000. Blots for CpRom1 were probed as described in Bertuccini et al. (2024). Blots for native CpRom2 and CpRom3 were probed with mouse pre-immune and immune sera diluted 1:250. Three washing steps of 5 min each were conducted with TNT buffer plus 0.1% BSA before the incubation with secondary antibodies. For all blots, incubation with secondary antibodies was conducted for 1 h, at 25°C, with goat anti-mouse IgG-HRP conjugate (Bio-Rad, Hercules, CA USA) diluted 1:3,000. Three washing steps as above were conducted after incubation with secondary antibodies. Detection of proteins was performed using Pierce ECL substrate (ThermoFisher), and bands were visualized with a ChemiDoc MP Imaging System (Bio-Rad, Hercules, CA, USA).

### Preparation of antisera for CpRom2 and CpRom3

Aliquots of purified and dialysed 6His-CpRom2 and 6His-CpRom3 were used to prepare mouse antisera as follows. Three Balb-C mice for each recombinant protein were immunized with 100 µg of protein plus complete Freund adjuvant as first inoculum: after 30 days, with 100 µg of protein plus incomplete Freund adjuvant as second inoculum, and, after another 30 days, with 100 µg of protein in PBS. Finally, mice were bled 15 days after the last inoculum and 150-200 µl of serum was obtained from each mouse. Mice sera were tested by dot blot spotting approximately 100 ng of recombinant protein on nitrocellulose membrane (Bio-Rad, USA) for spot in a Bio-Dot apparatus (Bio-Rad, USA), and spotted proteins were completely air dried with vacuum. Membranes were treated and developed as for the Western blots (see above).

### Fractioning of E. coli extract in M15, Ruv3 and Ruv 5 E. coli strains

Recombinant plasmids for expression of his-tagged rhomboids were also used to transform *E. coli* strains RV-1337-3 (RuV3) and RV-1337-5 (RuV5) (Massey-Gendel et al., 2009). For each transformed strain, 8 ml of LB medium was inoculated and let grow at 37 °C up to 0.6 OD, then 1 mM of IPTG was added to induce expression of recombinant rhomboids and let grow for additional 3 hours. Bacteria were pelleted by centrifugation at 2,040 x g, resuspended in 1 ml of H_2_O and subjected to 6 cycles of 30 sec of sonication at 75 % of strength in HD2070 Sonopuls (Bandelin Electronic, Berlin). Then, bacterial lysates were centrifuged at 10,000 x g for 15 min at 4 °C, pellets were resuspended in 50 μl of Laemmli buffer to analyse inclusion bodies composition and 10 μl were used for SDS-PAGE and Western blot analysis. Supernatants were centrifuged at 90,000 x g in Optima-MAX TL-ultracentrifuge (Beckman-Coulter, USA) to collect membrane fraction, resuspended in 50 μl of Laemmli buffer, and 10 μl were loaded in SDS-PAGE, while 50 μl of ultracentrifuge supernatants were diluted with 4 x Laemmli buffer and 12.5 μl loaded in SDS-PAGE as soluble protein fraction. Results are reported in Supplementary Figure S1 (supplementary data).

### Indirect immunofluorescence assay

Oocysts and sporozoites were fixed on microscopic slides with 4% formaldehyde in PBS for 10 min, washed, and blocked with 2% BSA in PBS for 1 h. Sporozoites were permeabilized with PBS plus 0.1 % Triton X-100 (Sigma-Aldrich) for 10 min at RT and then removed by washing the slides with PBS. In some experiments, permeabilization was omitted (i. e. Figure 10F-G). The anti-rhomboids mouse sera were diluted from 1:100 to 1:1000 in PBS plus 2% BSA and incubated for 1 h at room temperature. In the experiment in Figure 8 the anti-ID2 rabbit serum was used 1:1000 in PBS plus 2% BSA and incubated for 1 h at room temperature in combination with anti-CpRom1 (mouse serum) 1:1000. After washing in PBS, the slides were incubated for 1 h with goat anti-mouse antibody conjugated with fluorescein (FITC) (BioRad, Hercules, CA), diluted 1:1000 in PBS plus 2% BSA and with goat anti-rabbit antibody conjugated with rodamin 1:1000 (BioRad, Hercules, CA). All incubations were performed at room temperature. Finally, nuclei staining was obtained with DAPI (4ʹ,6-diamidino-2-phenylindole) 300 nM (Thermo Fisher) for 1 min at RT. Slides were then washed with PBS, and mounting medium (Thermo Fisher) was added before sealing the slides with the cover-slides.

Staining procedures for HCT8 infected cells were conducted as above on cells cultured on micro-slides chamber (Thermo Fisher) probed with mouse primary antibodies for rhomboids and rabbit anti-sporozoite serum 1:1000 (Trasarti et al., 2007). Mouse anti-rabbit Alexa (388)-conjugated antibody and goat anti-mouse Alexa (594)-conjugated antibody (Thermo Fisher) were used as secondary antibodies. Microscopic analysis was conducted with a fluorescence microscope (Zeiss-Axioplan 2), and images elaborated with Axiovision 4.8.1 (Zeiss, Germany).

### In vitro infection on HCT-8 cell culture

Human colorectal adenocarcinoma cells (HCT-8) were maintained in RPMI 1640 medium (GIBCO/Invitrogen, Paisley, United Kingdom), enriched with 5% fetal bovine serum (FBS; GIBCO), 200 mM L-glutamine (Sigma, St. Louis, MO), 1% sodium pyruvate (Sigma), and 5% penicillin and streptomycin. These cells were cultured in flasks and kept at 37 °C in a humidified atmosphere with 5% CO2 and 85% humidity. Aliquots of 1×10^7^ oocysts of *Cryptosporidium parvum* (Iowa strain) were induced to excyst (see above), and after 30 min the excystation mix was used to infect semi-confluent HCT8 cells. Infections were conducted with a multiplicity of infection of 10 oocysts for cell, and cultures were continued for 48 hours on chamber slides before fixing for microscopy.

### Ultrastructural analysis and immunolocalisation of CpRom1

For immunolocalization of Cprom1, samples containing free sporozoites were fixed overnight at 4 °C with 4% paraformaldehyde and 0.1% glutaraldehyde in 0.1 M Na cacodylate buffer. Next, samples were rinsed in 0.1 M Na cacodylate buffer, dehydrated in ethanol serial dilutions and embedded in LR White, medium-grade acrylic resin (London Resin Company, UK). Samples were polymerised in a 55 °C oven for 48 h, and ultrathin sections (90 nm) were collected on gold grids. The grids were floated on drops of PBS containing 0.1 M glycine for 10 min, washed with PBS, blocked with 5% normal goat serum/1% BSA in PBS for 30 min and incubated overnight at 4 °C with the anti-ID2 (1:10) in PBS/0.1% BSA. After washing, the grids were incubated for 1 h with 10 nm gold-conjugated goat anti-mouse IgG (Sigma Aldrich) diluted 1:20, rinsed in PBS/0.1% BSA, followed by distilled water and air dried. Finally, samples were stained successively with 2% uranyl acetate and Reynolds lead citrate solution, and observed with a Philips EM208S transmission electron microscope (FEI - Thermo Fisher).

### Bioinformatics tools

General similarity searches with DNA and protein sequences were conducted on non-redundant GenBank databases using the BLAST program, available at https://blast.ncbi.nlm.nih.gov/Blast.cgi. To identify protein coding sequences, ORFfinder at https://www.ncbi.nlm.nih.gov/orffinder/ and GenScan at http://genes.mit.edu/GENSCAN.html were used. Searches for homologous proteins were performed on databases of predicted proteins using Blastp algorithm at the following web sites: http://cryptodb.org/cryptodb/ for *Cryptosporidium* spp. and *Gregarina niphandrodes*. http://toxodb.org/toxo/ for *Toxoplasma gondii*; http://plasmodb.org/plasmo/ for *Plasmodium falciparum*;

The evolutionary history was inferred using the Neighbor-Joining method, bootstrap test was performed (500 replicates). The evolutionary distances were computed using the Poisson correction method and reported in the units of the number of amino acid substitutions per site. The analytical procedure encompassed 29 amino acid sequences. The pairwise deletion option was applied to all ambiguous positions for each sequence pair resulting in a final data set comprising 127 positions.

Evolutionary analyses were conducted in MEGA12 (Kumar S, et al., 2024) utilizing up to 4 parallel computing threads. Prediction of transmembrane domains (TMs) was performed with TMHMM v.2 at https://services.healthtech.dtu.dk/services/TMHMM-2.0/ and at http://smart.embl-heidelberg.deorwithDeepTMHMM-1.0 at https://services.healthtech.dtu.dk/services/DeepTMHMM-1.0/ (Hallgren et al., 2022). Newly predicted TMs were verified with AlphaFold (Fleming et al., 2025) at https://alphafold.ebi.ac.uk.

Clustal alignments were performed at http://www.ebi.ac.uk/Tools/msa/clustalo/. Bidimensional representations of CpRoms were obtained with Protter at http://wlab.ethz.ch/protter/start/ (Omasits et al., 2014), applying custom topology for the transmembrane domains. Putative targets of rhomboids were identified by similarity searches at CryptoDB with the following proteins of *Toxoplasma gondii*: TgMic2 (TGME49_201780), TgMic6 (TGME49_218520), TgMic12 (TGME49_267680), TgMic8 (TGME49_245490), and the *Plasmodium falciparum* thrombospondin-related anonymous protein (TRAP) (PF3D7_1335900.1) as queries in the BLAST program. Searches for protein-protein interaction were carried out in String database at https://string-db.org/cgi/input?sessionId=bV4mVyVVojwM&input_page_show_search=on.

## Results

### Identification of C. parvum rhomboids

CpRom1 identification was previously reported (Trasarti et al. 2007). In this study, other two predicted rhomboids were identified by a homology search conducted on the translated *C. parvum* (Iowa strain) genome, using various apicomplexan rhomboid sequences (i.e. known rhomboids from *T. gondii* and *P. falciparum*) as queries. Altogether, this parasite encodes for three functional rhomboids, and the assigned name, Crypto ID and UniProt ID are reported in Table 1. We noted that there was a clear case of homonymy regarding the CpRom1 name, given that this name was assigned to three different proteins as reported in Table 1 (Li et al., 2016; Castellanos-Gonzalez et al., 2019; Gao et al., 2021; Bertuccini et al., 2024). Therefore, the need for an unambiguous classification has prompted us to propose a revised nomenclature based on protein length, in decreasing order as reported in table 1. Hence, these proteins were named CpRom1 (990 aa), CpRom2 (464 aa) and CpRom3 (282 aa), and such nomenclature will be used throughout the manuscript. Genes encoding CpRom1 and CpRom2, namely cgd6_760 and cgd7_3020, respectively, are mono-exonic and do not contain introns. Differently, CpRom3, encoded by cgd3_980, is constituted by two exons (Figure 1).

**Figure 1.**
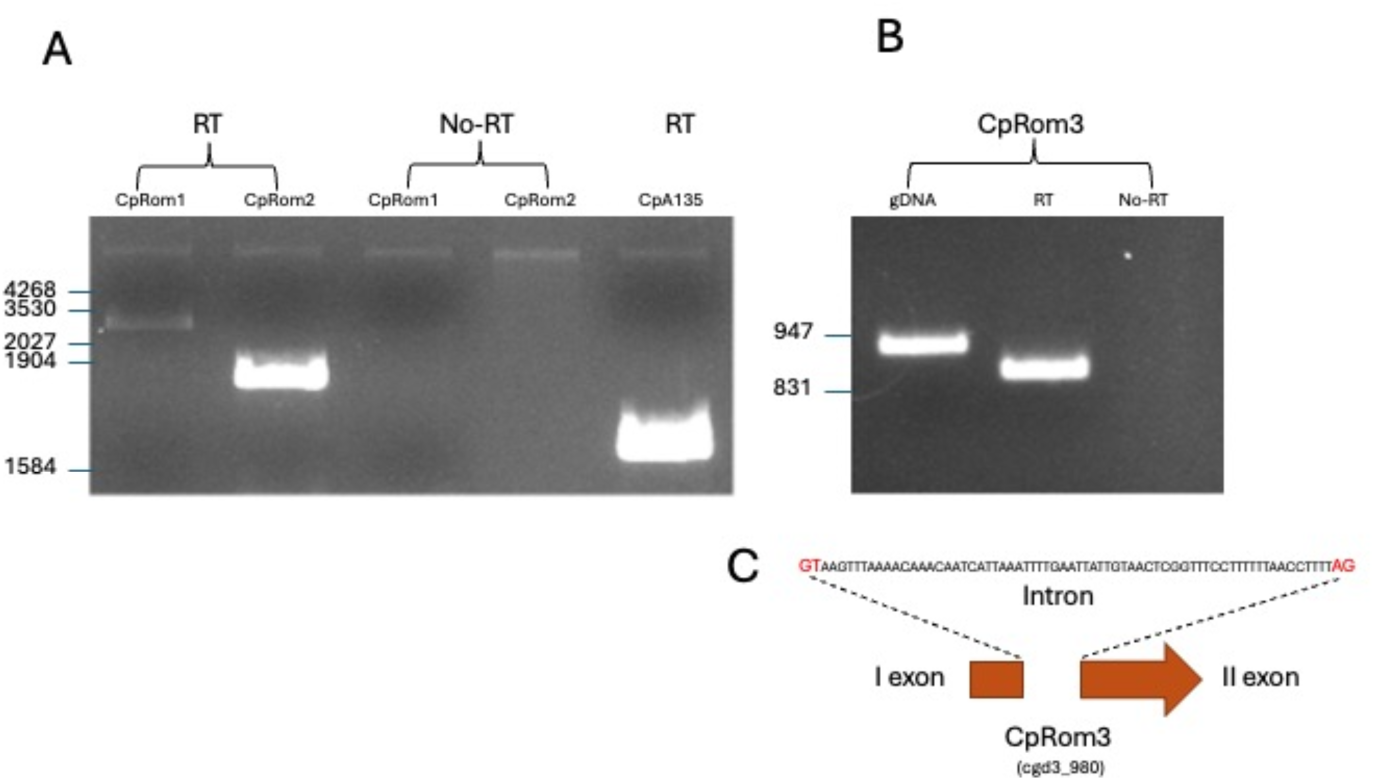
Expression of C. parvum rhomboids in sporozoites. A, RT-PCR on sporozoite mRNAs, 1-2 reverse transcribed mRNA and PCR for CpRom1 (lane 1) and CpRom2 (lane 2), 3-4 PCR for CpRom1 (lane 3) and CpRom2 (lane 4) on non-reverse transcribed mRNA (negative controls). Lane 5, PCR amplification for CpA135 on reverse transcribed mRNA (Tosini F et al., 2004). B, 1-2 PCR amplification for CpRom3 on genomic DNA (lane 1) and on reverse transcribed sporozoite mRNAs (lane 2) respectively; 3, PCR amplification for CpRom3 on non-reverse transcribed mRNAs (negative control). 3, scheme showing the two exons encoding CpRom3 and the interposed intron sequence.

### Expression analysis of CpRom genes in sporozoites

To ascertain the expression of the three CpRom genes, three couples of primers were designed based on their genomic sequences and used for RT-PCR experiments on total RNA extracted from excysted sporozoites. As shown in Figure 1, all these genes are expressed in the sporozoites. The RT-PCR experiment also confirmed that the CpRom3 gene (Figure 1B-C) is composed of two exons and one intron, and its sequence is reported in Figure 1C.

### Phylogenetic Comparison of C. parvum rhomboids with the other apicomplexan rhomboids

The *C. parvum* rhomboids identified were then compared with predicted proteins of other apicomplexan parasites in a genome-wide search to produce a representative phylogenetic tree of Apicomplexa rhomboids (Figure 2). This analysis was conducted on the genomes of reference strains fully sequenced and annotated in EuPathDB, and *Cryptosporidium muris*, which is also annotated in CryptoDB, was also included as a representative of the *Cryptosporidium* species infecting the stomach instead of the intestine. The phylogenetic tree in Figure 2 shows that CpRom1 and CpRom2 (in red in the tree) are strictly related to PfRom4, TgRom4 and TgRom5 involved in cleaving the micronemal adhesins (Brossier et al. 2005; Baker et al., 2006; Rugarabamu et al., 2015). Differently, CpRom3 (also in red in the tree) appears to be related to a different cluster that includes PfRom3 and TgRom3. Genomic comparison also showed that none of the *C. parvum* rhomboids could be assigned to the PARLs cluster. Of note, the strictly related *C*. *muris* showed an additional fourth rhomboid that we named CmRom4 and that clustered with the PARL-like rhomboids (see also below).

**Figure 2.**
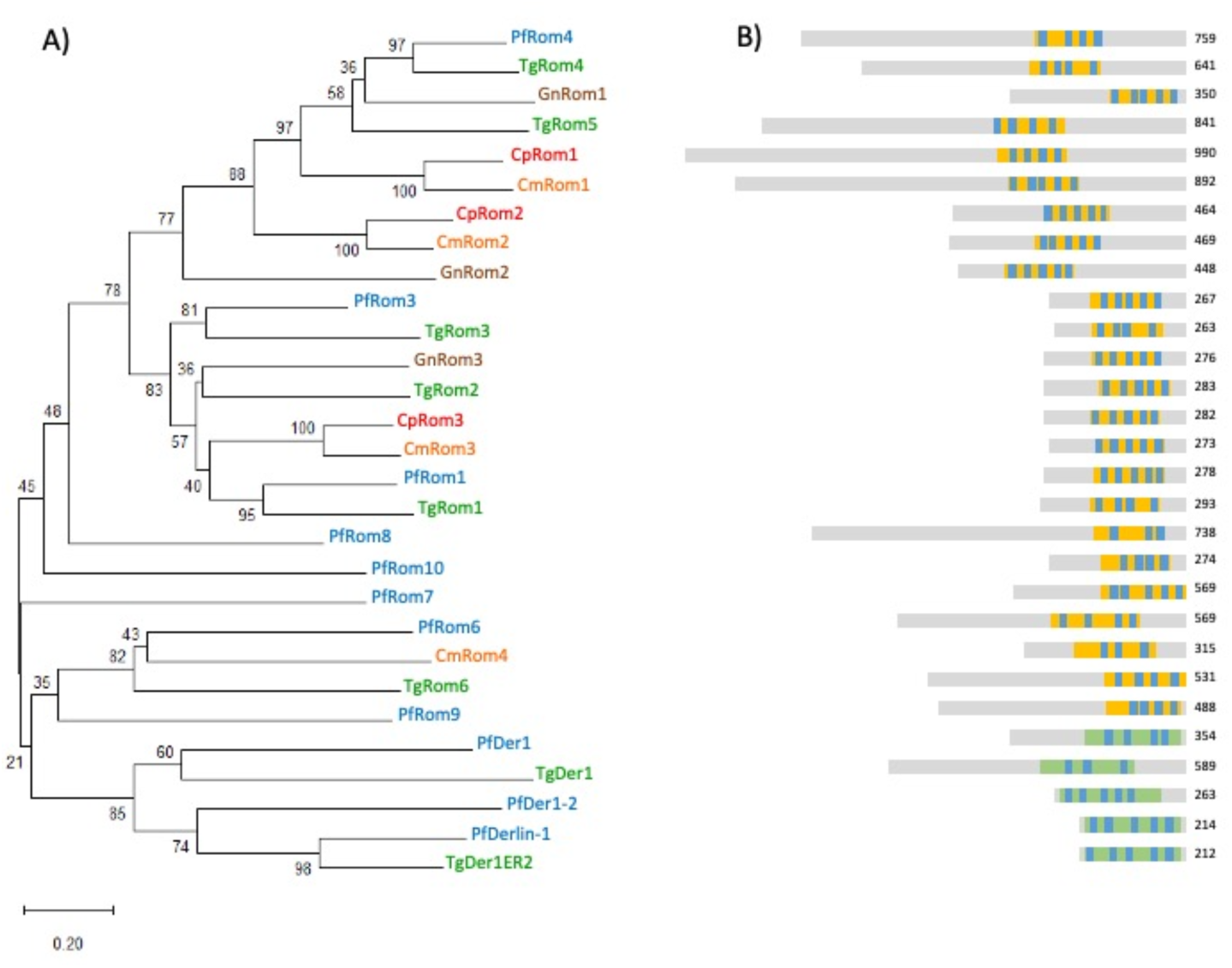
Evolutionary relationships of Apicomplexa rhomboids. A) Neighbor-Joining phylogenetic tree of the Apicomplexa rhomboids and rhomboid-related proteins (Derlins) is shown. The percentage of replicate trees in which the associated taxa clustered together in the bootstrap test are shown next to the branches. The tree is drawn to scale, with branch lengths in the same units as those of the evolutionary distances used to infer the phylogenetic tree. Protein names are coloured according to species. B) A schematic representation of proteins is proposed, the rhomboid domain is reported as yellow bars while Derlin domain as green bars. Transmembrane helices are indicated by blue segments (list of TMs with their position are reported in a Supplementary file S2). The lengths of the proteins are indicated on the right.

#### Identification of structural domains and catalytic sites of C. parvum rhomboids

To identify structural domains and the catalytic sites along the aminoacidic sequences of the three *C. parvum* rhomboids, their sequences were aligned with those of the most similar rhomboids from *P. falciparum* and *T. gondii* (see Figure 2), which are the most characterized among the apicomplexan rhomboids. These alignments led to the identification of further TM domains that were not previously identified and were confirmed by AlphaFold modelling. Novel TM domains were revealed in TgRom1, TgRom4, TgRom5, PfRom4 as well as in CpRom1 and are highlighted in Figure 3. Moreover, the alignment allowed us to determine the exact positions of the critical aa that compose the catalytic sites and that are distributed between TM4 and TM6. The results of this analysis are shown in Figure 3.

**Figure 3.**
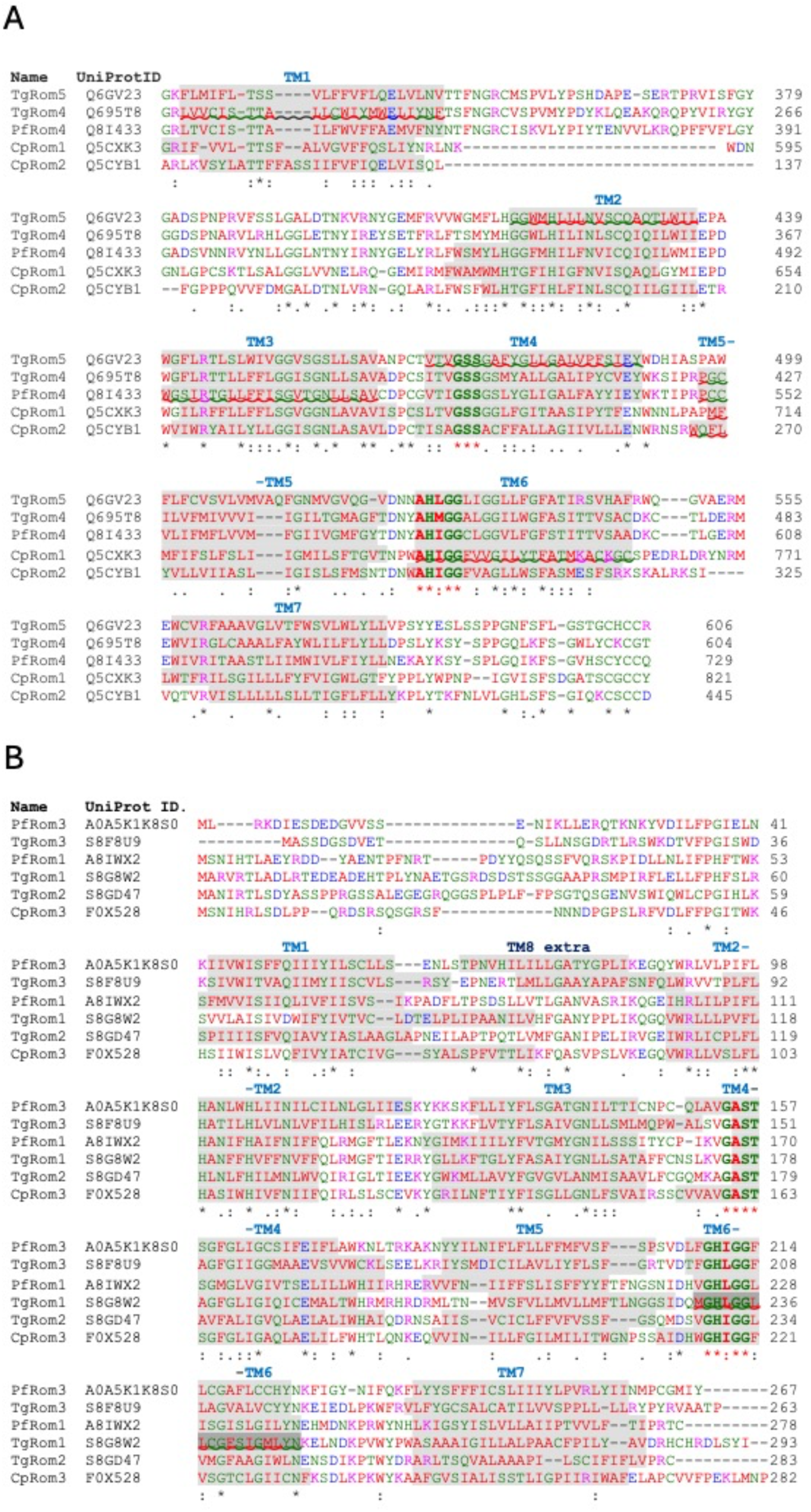
A, Alignments with homologs of CpRom1 and CpRom2 showing TM domains and catalytic sites. Alignments were obtained with ClustalO. Gross part of the proteins at their N-terminus were eliminated to align the rhomboid domains of the proteins. B, TM domains and catalytic sites in CpRom3 aligned with similar proteins of Toxoplasma gondii and Plasmodium falciparum. Alignments were obtained with ClustalO. Shaded amino acids indicate predicted TM domains. Waved underlines indicate predicted TM domains obtained directly with TMHMM v.2. Wavy underscored TM domains were predicted based on the most similar overlapping sequence and verified by AlphaFold. Bold letters correspond to the catalytic dyad and the surrounding conserved amino acids. FPHF in blue dotted box indicate post-Golgi-sorting motif and FF in green dotted boxes indicate Golgi-targeting motif (Sheiner L et al., 2008).

As expected, CpRom1 and CpRom2 clustered with TgRom4, TgRom5 and PfRom4, all sharing the following conserved consensuses: a GSS motif at the fourth TM and an AHXGG motif at the sixth TM (Figure 3A). Differently, CpRom3 belonged to a separate lineage, which included PfRom3, TgRom3, PfRom1, TgRom1 and TgRom2. This group was characterized by the following conserved consensuses: a GAST motif at the fourth TM and a GHIGG consensus at the sixth TM. Furthermore, four rhomboids of this second group have a Golgi translocation signal before the TM (Figure 3B), namely the two consecutive phenylalanine (FF) in CpRom3 and TgRom2 and the phenylalanine-proline-histidine-phenylalanine (FPHF) motif in TgRom1 and PfRom1 (Sheiner et al., 2008).

A bidimensional representation of the three *C. parvum* rhomboids with the dislocation of the TM domains and the catalytic sites in relation to the cell membrane is depicted in Figure 4. Finally, these results showed that the active apicomplexan rhomboids, except for PARLs, are composed of 7 TM domains and can be classified as mixed-secretase (Lemberg and Freeman, 2007). Furthermore, these apicomplexan rhomboids can be sub-grouped into two different clusters based on the conserved residues around the catalytic dyad.

**Figure 4.**
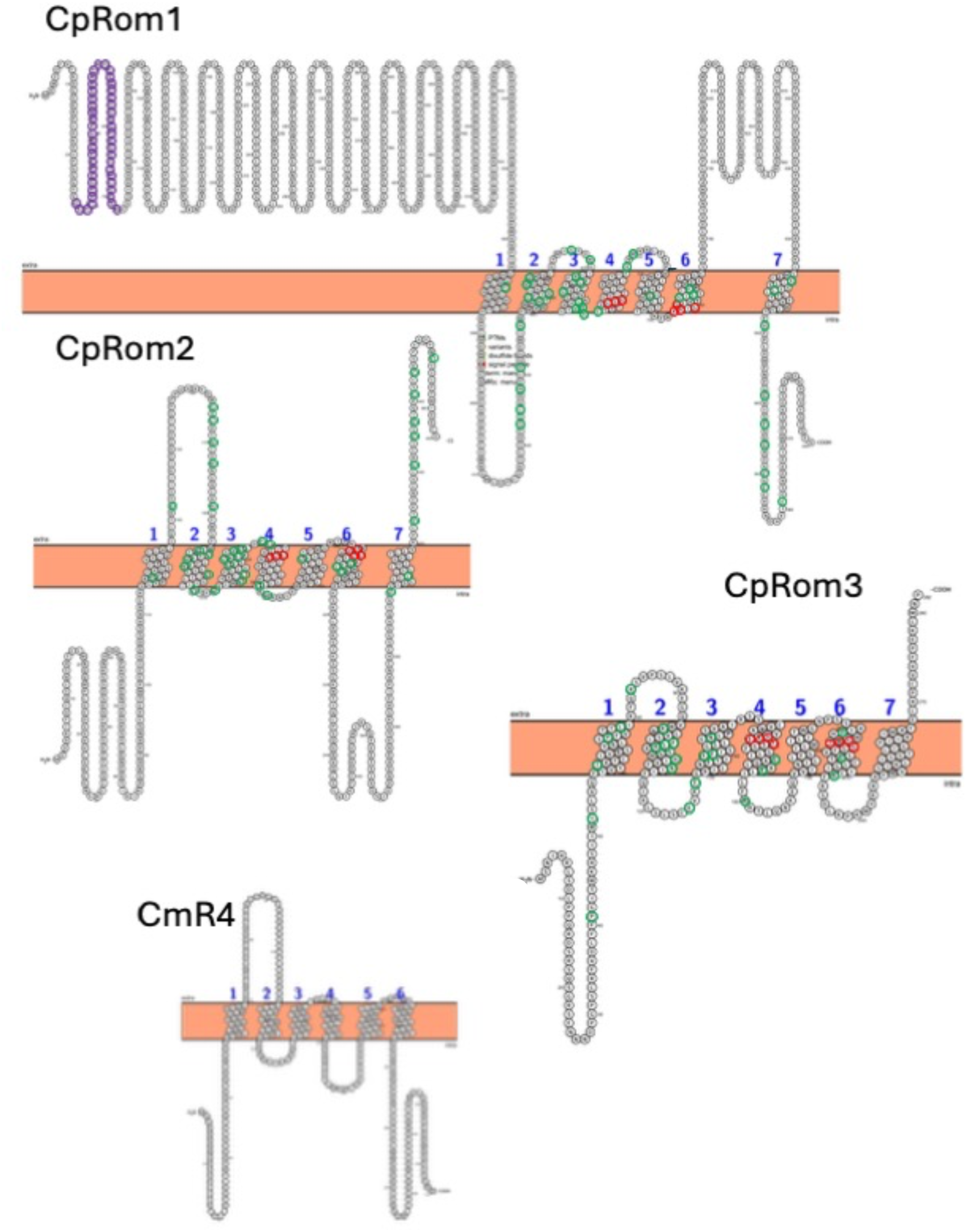
Bidimensional representation of the three *C. parvum* rhomboids and the PARL-like rhomboid CmR4 from *Cryptosporidium muris*. The circled amino acids represent the residues always conserved in the two clusters, namely one for CpRom1 and CpRom2 and the second for CpRom3, in red for the catalytic site and in green along the protein sequence (see in the text).

### PARL rhomboids in Cryptosporidium genus

A survey to identify rhomboids in genomes of other *Cryptosporidium* species revealed that species affecting the intestine (i. e. *C. hominis*, *Cryptosporidium meleagridis*) showed a similar genomic content with only three rhomboids (data not shown), differently two species, namely *C. muris* and *Cryptosporidium andersoni*, which infect the stomach encode for four rhomboids. None of the *C. parvum* rhomboids belong to the PARL group of mitochondrial rhomboids and this is not surprising given that this parasite as well as other *Cryptosporidium* species lack a functional mitochondrion. However, the similarity search for rhomboids in the Apicomplexa genus revealed that *C. muris* and *C. andersoni* both possess PARL-like genes. The protein CmRom4, the PARL-like rhomboid of *C. muris*, was more like the PARL proteins of *P. falciparum* and *T. gondii* (see Figure 2) than the other *C. parvum* rhomboids. A bidimensional representation of CmRom4 PARL-like rhomboid is reported in Figure 4B and, differently from the *C. parvum* rhomboids (Figure 4) shows only 6 TM domains. Overall, we found out that species that infect the intestine like *C. parvum*, *C. hominis*, *C. bovis* lack a PARL-like gene. By contrast, Cryptosporidium species infecting the stomach, namely *C. muris* and *C. andersoni*, possess a PARL-like rhomboid. There is no evidence for the presence of a mitochondrion in *C. andersoni*, although this species possesses some genes involved in the aerobic metabolism that are absent in *C. parvum* and *C. hominis* (Liu et al., 2016). Importantly, a mitochondrion has been identified at the ultrastructural level in *C. muris* (Uni et al., 1987). The genomic regions surrounding the PARL-like genes in *Cryptosporidium* species, by comparison, revealed a precise removal of these genes in species without mitochondrion like *C. parvum*, *C. hominis*, *C. bovis* and others (Figure 5). Remarkably, except for the precise excision of the PARL-like genes, the synteny, namely the distribution of the other surrounding genes along the chromosome, is perfectly conserved among these different species.

**Figure 5.**
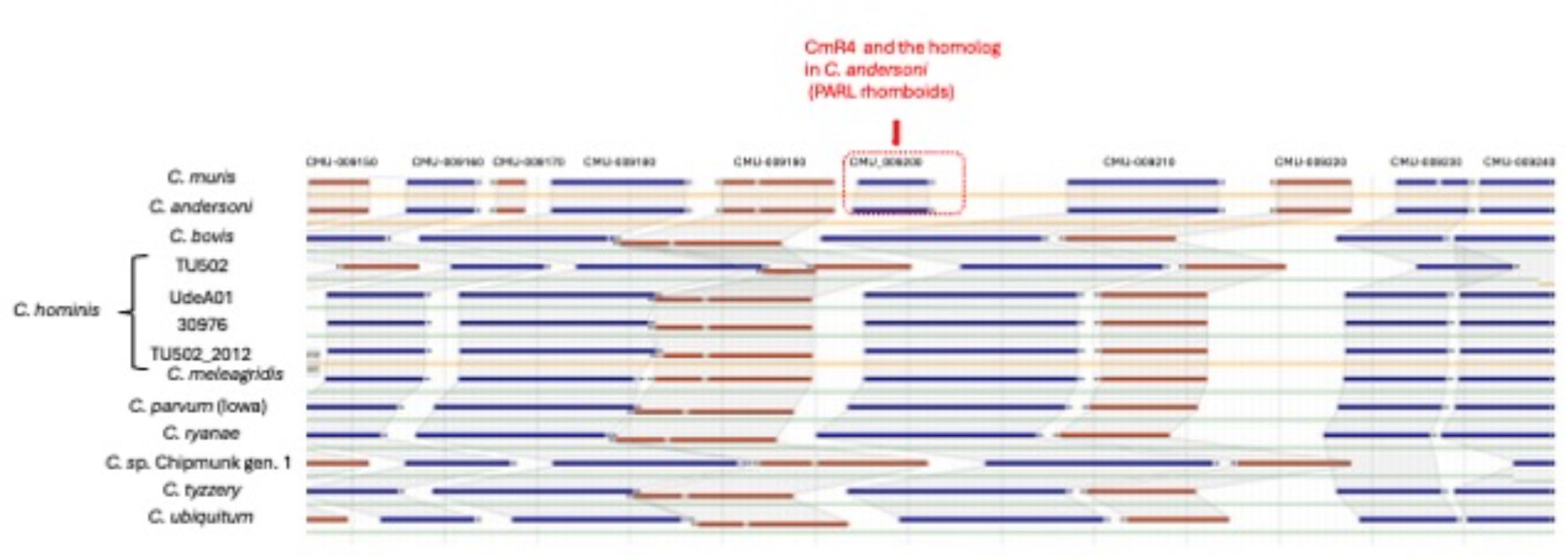
Genomic region of *Cryptosporidium muris* CMU_009200 gene with the surrounding genes and the syntenic genomic regions in other *Cryptosporidium* species as represented in CryptoDB. CMU_009200 gene encodes for the CmR4, the mitochondrial PARL rhomboid of this species, and the dotted red box highlights the two PARLs in the “gastric species” *Cryptosporidium muris* and *Cryptosporidium andersoni* that lack in the other “intestinal species”. Shaded areas embrace orthologs in the different species.

### Expression of recombinant C. parvum rhomboids in Escherichia coli

Coding sequences for the three rhomboids were cloned in expression vectors to produce recombinant proteins fused with a histidine tag. Portions of the coding sequences avoiding the multi-spanning rhomboid domain were also cloned, so to favour the expression of soluble portions of the proteins to obtain specific peptides for the immunization. Surprisingly, only the full-length tagged proteins, but not their shorter fragments, were expressed by bacteria. Schematic representations of the whole set of constructs and their expression results are reported (Supplementary Figure S2). Therefore, the full-length recombinant rhomboids produced were named 6h-CpRom1 (Bertuccini et al., 2024), 6h-CpRom2 and 6h-CpRom3, and their expected molecular masses were 110 kDa, 51 kDa and 35 kDa, respectively. However, the expression in a conventional recipient *E. coli* strain for his-tagged constructs (M15 strain) gave fragmented and scarcely soluble products that required the presence of denaturing agents (i. e. guanidine and urea) for the purification of the recombinant peptides (Figure 6). Thus, the recombinant 6h-CpRom2 and 6h-CpRom3 were synthetized as unique peptides of 20 kDa and 25 kDa respectively. The cutting of these recombinant proteins was probably due to the action of bacterial proteases before the lysis in denaturing conditions. In any case, the shorter 6h-CpRom2 and 6h-CpRom3 were suitable for the chromatographic purification and the preparation of specific mouse antisera. The successful production of specific IgGs was checked by dot blots at 60-and 75-days post immunization (Supplementary Figure S3).

**Figure 6.**
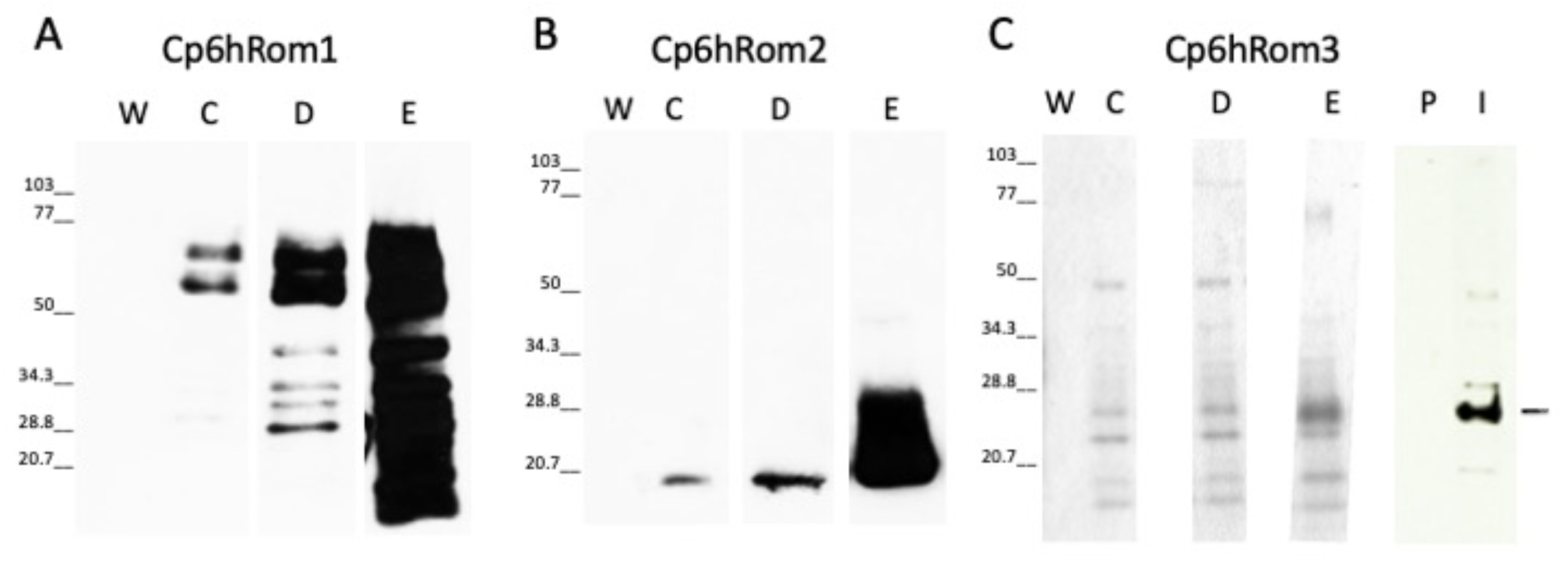
A, B, Western blot experiments the recombinant C. parvum rhomboids with 6 histidine tags at their N-terminus, the anti-tag (RGS-6his) MoAb (Qiagen) was used to detect the recombinant products. C, SDS-PAGE and Western blot with recombinant rhomboids. Lanes W-E: SDS-PAGE with the steps of chromatographic purification of recombinant CpRom3. The recombinant CpRom3 was scarcely detected by RGS-6his MoAb during the purification steps, despite is detected in the crude bacterial lysate (see Supplementary Figure S1). Lanes P-I: Western blot with pre-immune (P) and immune (I) mouse serum for CpRom3.

To produce functional recombinant rhomboids localized in cell membrane the expression of these constructs as well as that of 6h-CpRom1 was attempted in alternative *E. coli* strains which have been improved to produce functional membrane proteins (Massey-Gendel et al. 2009). Therefore, these strains were transformed with the 6h-CpRom1, 6h-CpRom2 and 6h-CpRom3 constructs and induced to express the recombinant rhomboids. The induced bacterial cultures were lysate, homogenized and separate in subcellular fractions that were analysed by Western blot (Supplementary Figure S1). None of the assayed recipient strain was compatible with the proper expression of 6h-CpRom1 and this recombinant protein was always accumulated in cytoplasmic inclusion bodies regardless of the strain. In Figure S2 the impaired expression in Ruv3 is shown and similar results were obtained with the other strains (data not shown). On the other hand, the expected bands of 51 and 35 kDa for 6h-CpRom2 and 6h-CpRom3 respectively were identified in the membrane fraction of Ruv5 strain. Therefore, the placement in a cell membrane was required to obtain the integral forms of these two proteins.

### Identification of C. parvum rhomboids in oocyst and sporozoite proteomes

To identify the native *C. parvum* rhomboids, whole sporozoite lysates at 0, 30 and 60 minutes after the start of the excystation were analysed by western blot using positive sera (Figure 7). Using anti-CpRom2 serum, we observed a unique band of approximately 70 kDa of constant intensity (Figure 7A) from the oocyst (time 0) to the excysted sporozoites (time 30 and 60 min from the excystation start). The difference between the expected size of approximately 50 kDa and the observed size of 70 kDa can be the result of a post-translational modification such as the addition of glycosides or other covalently linked molecules. Differently, the native CpRom3 appeared as a triplet as revealed by the anti-CpRom3 serum: one of approximately 70 kDa and two smaller bands of approximately 30 and 35 kDa that vary in intensity from the beginning up to one hour from the excystation start (Figure 7B). The 35 kDa band corresponded exactly to the expected size of CpRom3, appeared only after the excystation and its amount slightly increased in one hour after the excystation (Figure 7B). In contrast, the 30 kDa form decreased during the excystation (Figure 7B). The presence of three forms of this protein can be explained by processes that modify CpRom3 in the sporozoite cell. Some rhomboids, by paradox, can assume a dimeric form in SDS-PAGE (Kreutzberger and Urban, 2018), and this may explain the 70 kDa form of CpRom3, which is exactly the double of the expected 35 kDa.

**Figure 7.**
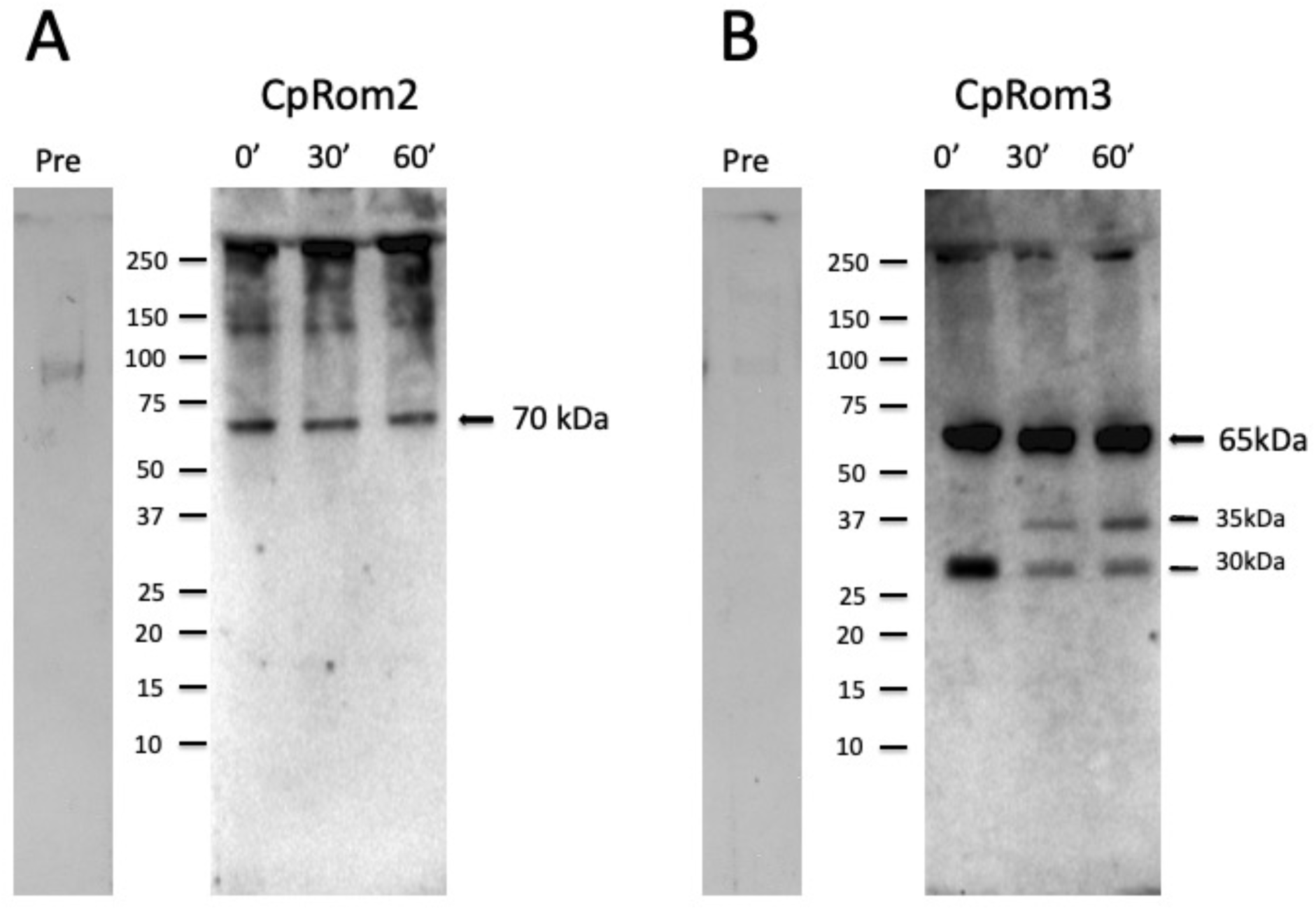
Western blot analysis of oocyst and sporozoite lysates at different times from the induction of excystation. A, lysates were probed with polyclonal mouse anti-serum for CpRom2 (dilution 1:500). B, lysates were probed with polyclonal mouse anti-serum for CpRom3 (dilution 1:100). Numerals at the top indicate minutes from the excystation start. Lanes marked with Pre show Western blots probed with pre-immunised mouse sera respectively for CpRom2 (left) and for CpRom3 (right). Molecular standards are reported on the left side of the blots.

**Figure 8.**
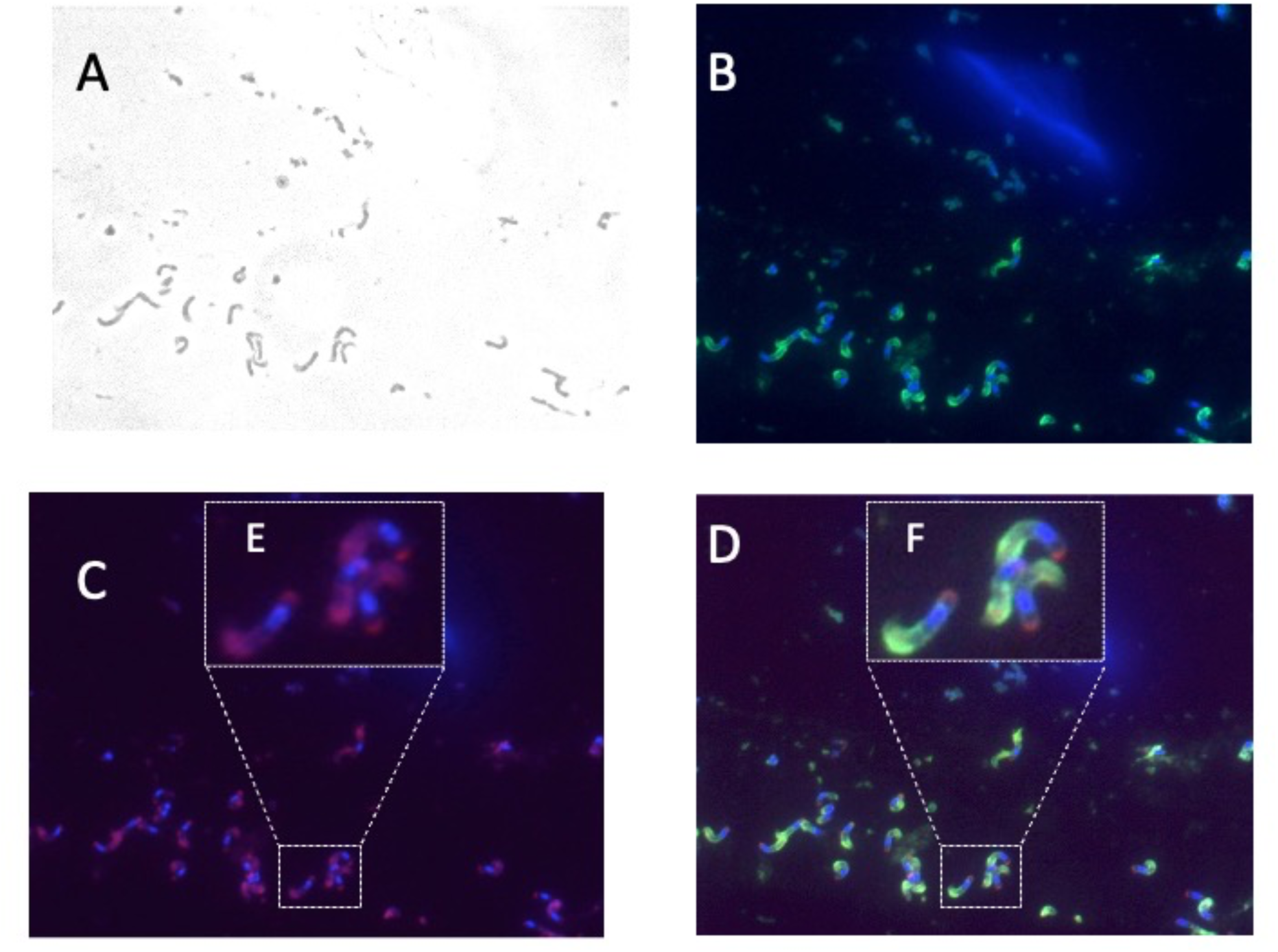
Localization of CpRom1 in excysted sporozoites (1000 X). A, excysted sporozoites in white field. B, sporozoites labelled with DAPI (blue) and anti-sporozoite serum (green). C, anti-CpRom1 serum. D, merge of B and D. E and F, enlarged portions of C and D respectively (dotted rectangles) to highlight labelling at the anterior and posterior pole of sporozoites.

Overall, the three *C. parvum* rhomboids, including the CpRom1 previously described (Bertuccini et al., 2024), are present in the oocyst and expressed in the excysted sporozoites.

### Different localization of C. parvum rhomboids in sporozoite

The specific sera for *C. parvum* rhomboids were also used to localize these proteins and the colocalization with the nucleus marked with DAPI was used for placing the rhomboids respect to the anteroposterior axis of sporozoites.

For CpRom1, we also used a rabbit antiserum for sporozoite antigens that intensely stained the apical surface of sporozoites and the parasitophore vacuole (Tosini et al., 2019). Therefore, the combination of nucleus (blue) and the apical surface (green) labelling revealed that the CpRom1 had a dual location: one apical, evenly distributed and, apparently, inside the cell membrane; a second accumulation point that was instead exactly at the posterior pole of the sporozoites (Figure 8). The immune electron-microscopy experiments confirmed the dual distribution of CpRom1 at the apical and posterior poles of sporozoites, as well as its internal localization at the apical pole (Figure 9).

**Figure 9.**
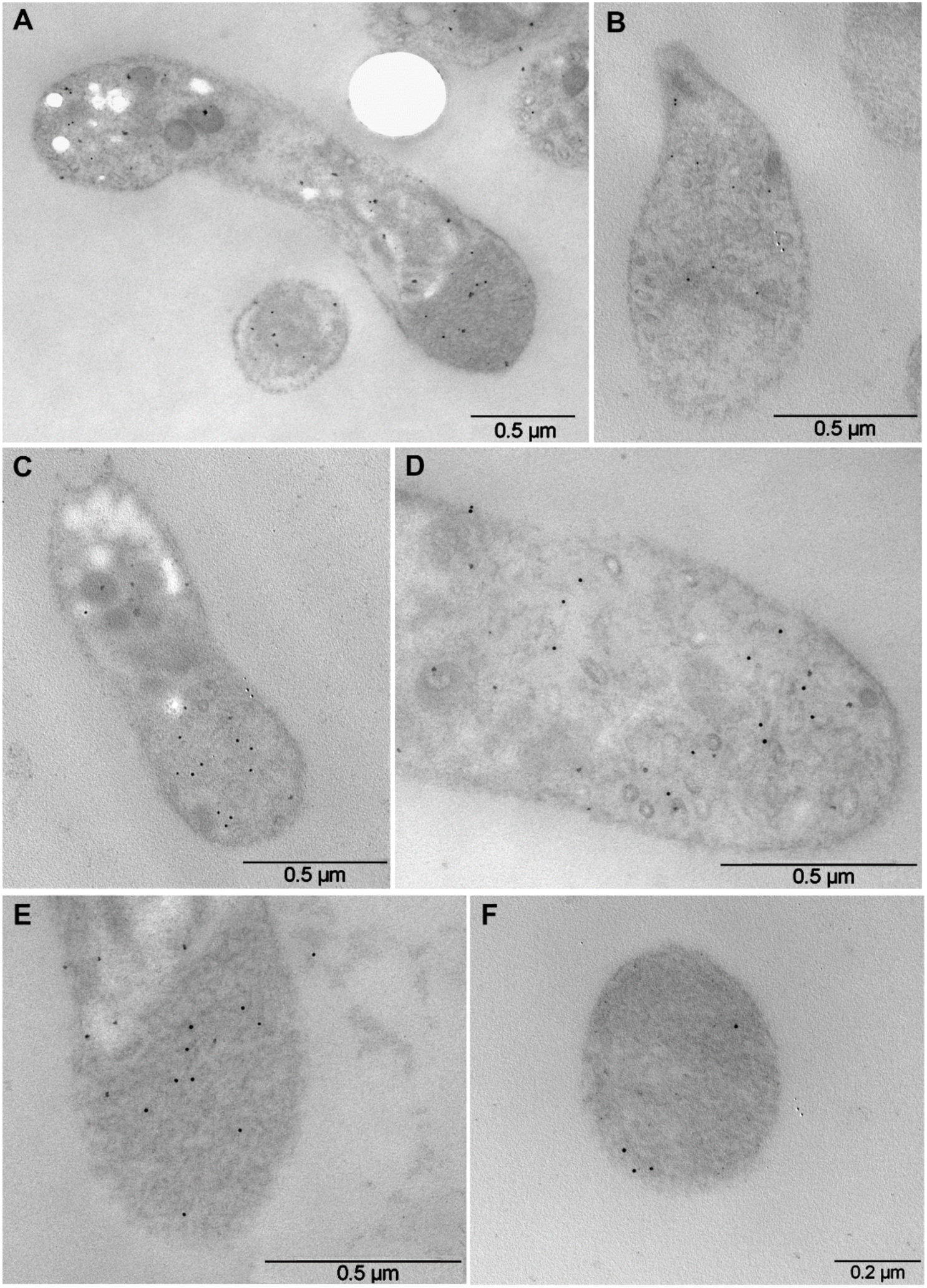
Immunoelectron microscopy of Cryptosporidium sporozoite ultrathin sections showing the dual localization of CpRom1 obtained by rabbit IgG for the ID2 antigenic portion. A, longitudinal section of an entire sporozoite showing the immunogold particles at the apical and posterior pole. B, longitudinal section of the apical complex with micronemes. C, an area of the micronemes adjacent to the dense granules. D, a high magnification view of the apical pole with rhoptry. E, longitudinal sections at the posterior pole at the level of the crystalloid body. F, a cross-section of the crystalloid body.

**Figure 10.**
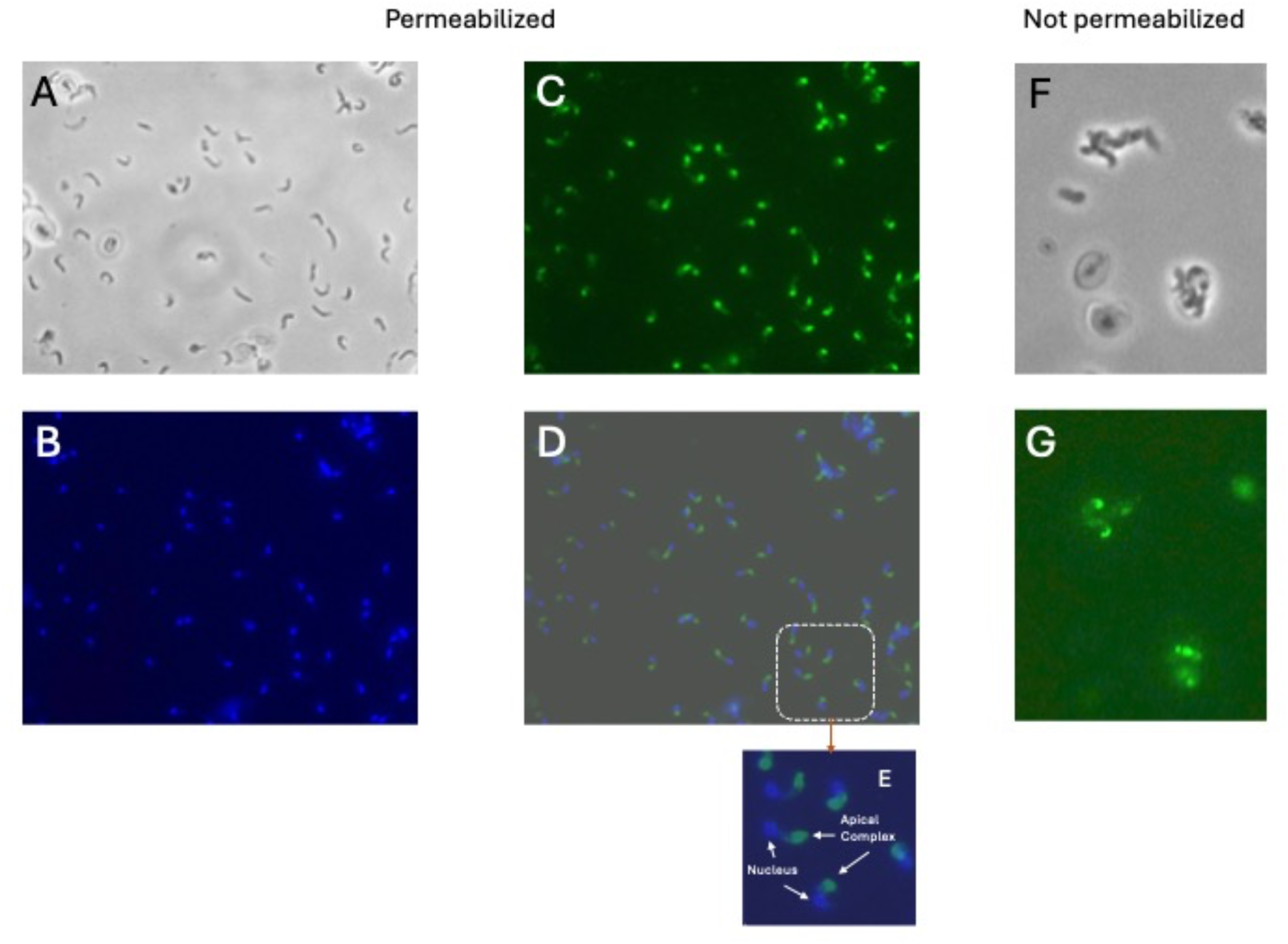
Immunolocalization of CpRom2 in excysted sporozoites (630 X). A-E, Colocalization on permeabilized excysted sporozoites. A, excysted sporozoites in white field. B, sporozoites labelled with DAPI (blue) and anti-sporozoite serum (green). C, anti-CpRom2 serum. D, merge of B and C. E, digital magnification of white dotted rectangle in D. F-G, imuunolocalization of CpRom2 on not permeabilized excysted sporozoites. F, excysted sporozoites in white field. G, immunofluorescence on the same field anti-CpRom2 serum.

CpRom2 was instead distributed exclusively in the anterior portion of the sporozoite around the apical complex (Figure 10). This localization was evident in the digital magnification of few sporozoites in Figure 10E. We also tried to place this rhomboid in relation to the external membrane repeating this immunolocalization on not permeabilized sporozoites and, as shown in Figures 10F and 10G, only some sporozoites (i. e. approximately two out six sporozoites within 30 min from the excystation start in this experiment) could be labelled in the upper part of the apical end. This resultsuggested that this rhomboid is accumulated in some apical organelle, may be the micronemes, and progressively transported to the apex of sporozoite.

Differently, CpRom3 was observed to be evenly distributed along the sporozoite surface (Figure 11), with a thickening exactly at the apical end of the sporozoite as shown with a digital magnification (Figure 11E).

**Figure 11.**
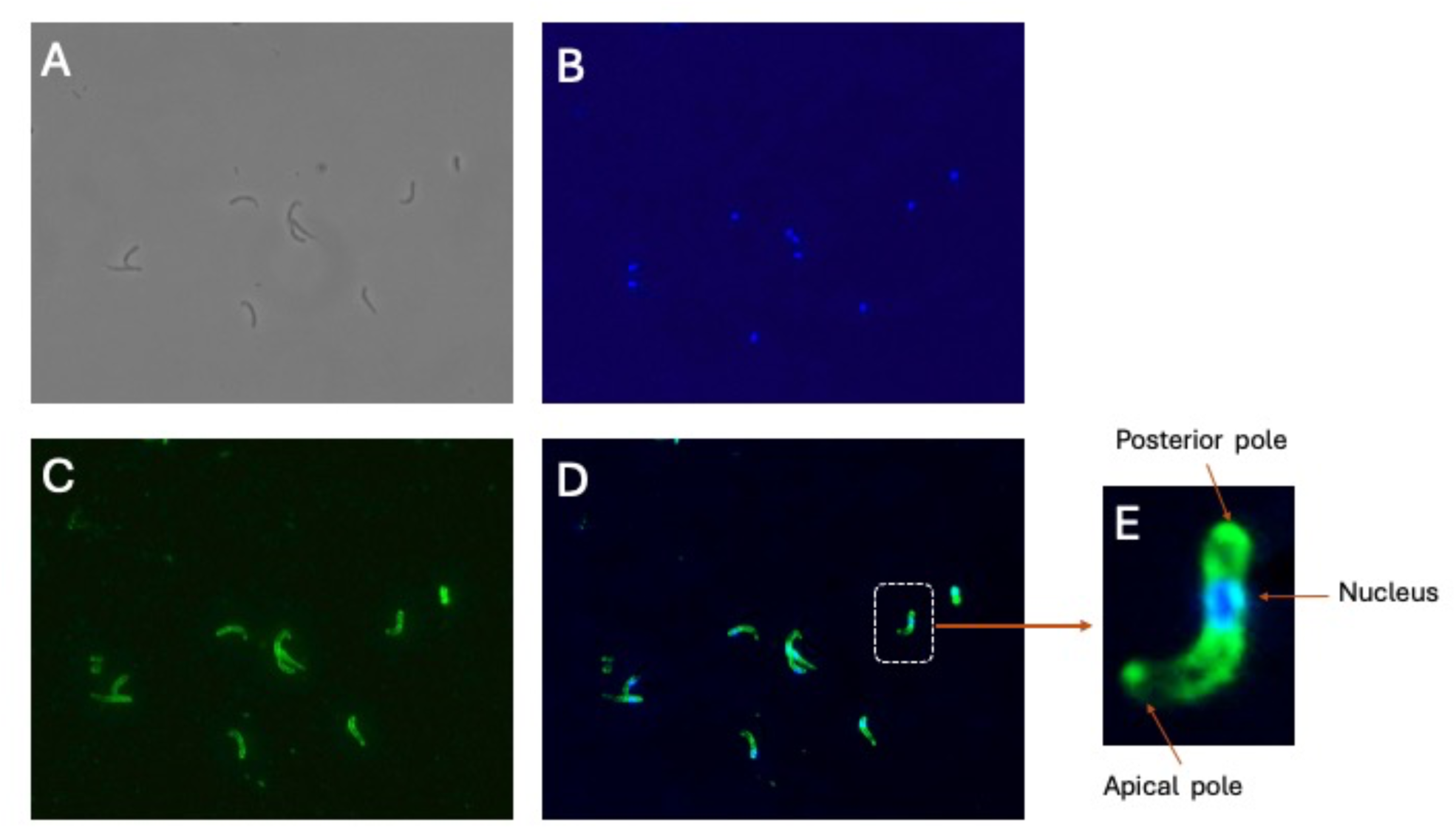
Localization of CpRom3 in excysted sporozoites (1000 X). A, excysted sporozoites in white field. B, sporozoites’ nuclei labelled with DAPI (blue). C, anti-CpRom3 serum. D, merge of B and C images. E, digital magnification of the sporozoite in the white dotted rectangle in D to localize the distribution of fluorescence along the sporozoite.

**Figure 12.**
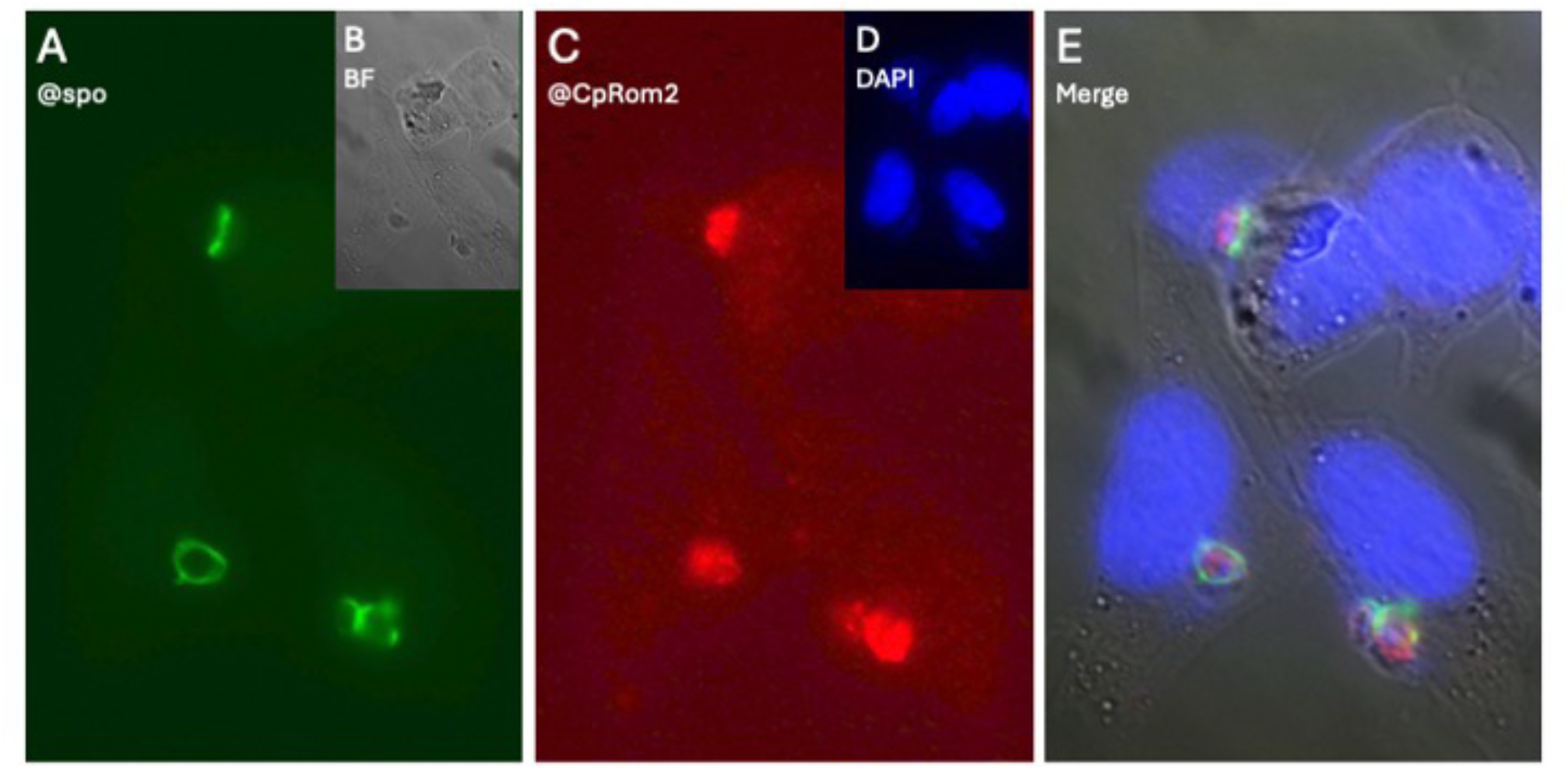
Colocalization of CpRom2 in infected HCT8 cells at 48 hours PI. A, parasitophorous vacuoles at different stages labelled with antisporozoite rabbit serum and mouse anti-rabbit Alexa (388)-conjugated antibody. B, microscopic field observed with white light (BF = bright field). C, parasitophorous vacuoles labelled with mouse anti-CpRom2 and goat anti-mouse Alexa (594)-conjugated antibody. D, DAPI staining of the nuclei. E, digital merging of A, B, C and D images.

Finally, these results clearly indicated that the three *C. parvum* rhomboids were coexisting in excysted sporozoites, although their spatial localization was different one from the other along the sporozoite cell.

### CpRom2 is present in the parasitophore vacuole in first stage of the infection

To detect rhomboids in the intracellular stages of the parasite in the first 48 hours post-infection, we also tested HCT8 infected cells by immunofluorescence. While we did not observe specific labelling for CpRom1 (Supplementary Figure S4) and CpRom3 (data not shown), CpRom2 was still present in the parasitophorous vacuoles (PVs) at different intracellular stages (Figure 11). In fact, the rabbit antiserum labelled in brilliant green the structure of PV (Tosini et al., 2009) and the mouse antiserum for CpRom2 marked in red the parasites inside the PV.

### In silico identification of substrates for C. parvum rhomboids

The cutting sites of rhomboids are constituted by few amino acids in the TM domain of the substrate membrane proteins (Strisovsky et al., 2009). However, a similarity search using the short consensuses of various proven rhomboid substrates in *C. parvum* genome did not reveal any consistent homology among the membrane proteins of this parasite. Therefore, the search for the putative targets of the three *C. parvum* rhomboids was based on the following criteria: high similarity with other apicomplexan proven substrates; the presence of one or more well recognizable TM domains; the co-expression with rhomboids in the sporozoite stage and, possibly, the colocalization with a rhomboid in a subcellular structure.

The similarity search on the *C. parvum* genome used the following substrates of apicomplexan rhomboids (Zhou et al., 2004): TgMic2 (TGME49_201780), TgMic6 (TGME49_218520), TgMic12 (TGME49_267680), TgMic8 (TGME49_245490) of *T. gondii* and the thrombospondin-related anonymous protein (PF3D7_1335900) of *P. falciparum.* These proteins share a micronemal localization and a modular architecture that implies repeated thrombospondin domains and/or EGF domains, which are involved in the progression toward and the attachment to the host cell, and the removal from the parasite surface mediated by a rhomboid cleavage. This first step revealed a total of 18 *C. parvum* proteins, which resulted very similar to two or more template proteins of *T. gondii* and *P. falciparum*. *C. parvum* proteins without an identifiable TM domain were eliminated from this group and only 10 proteins remained candidates as rhomboid substrates (Table 2).

**Table 2.**
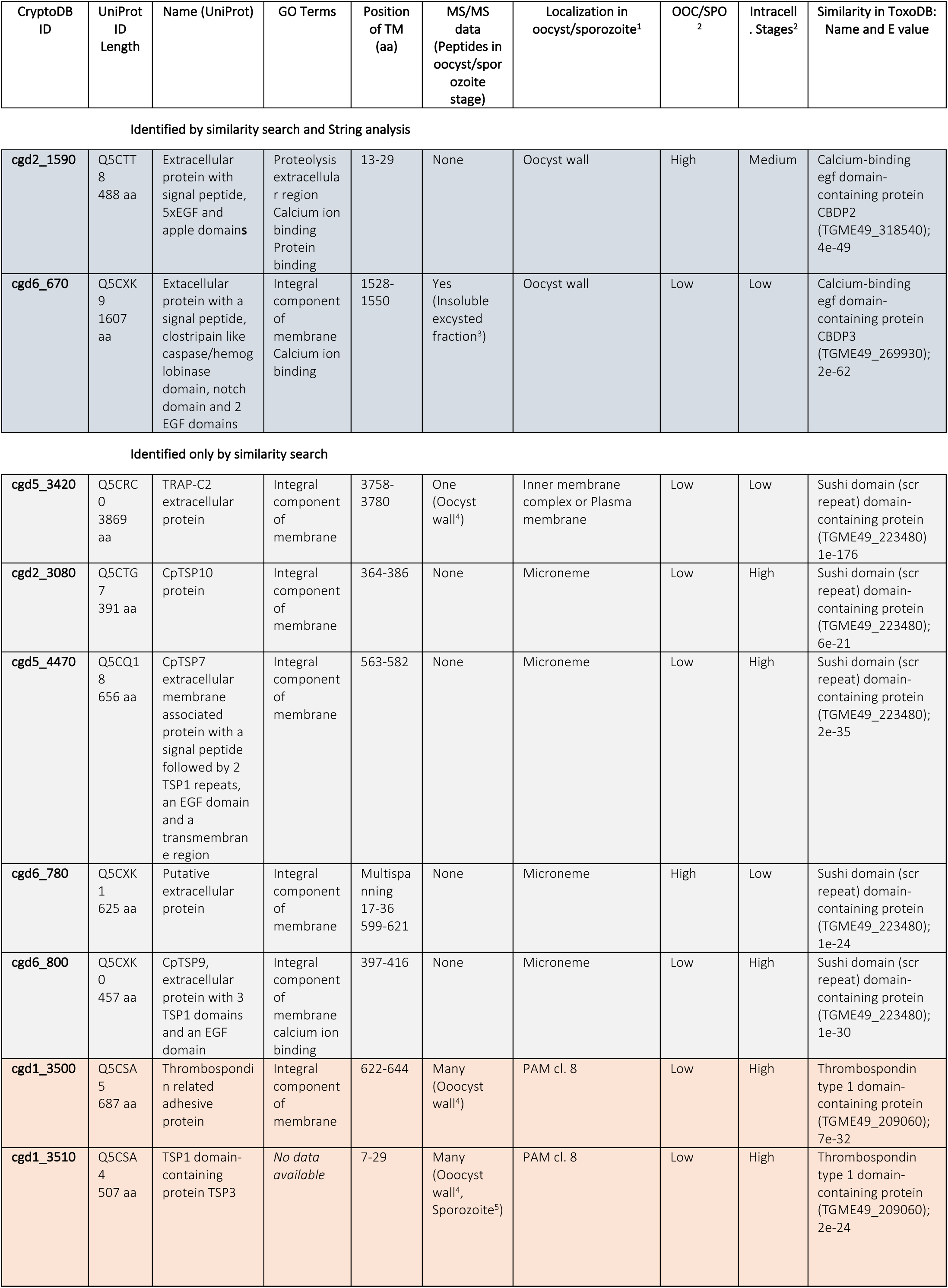
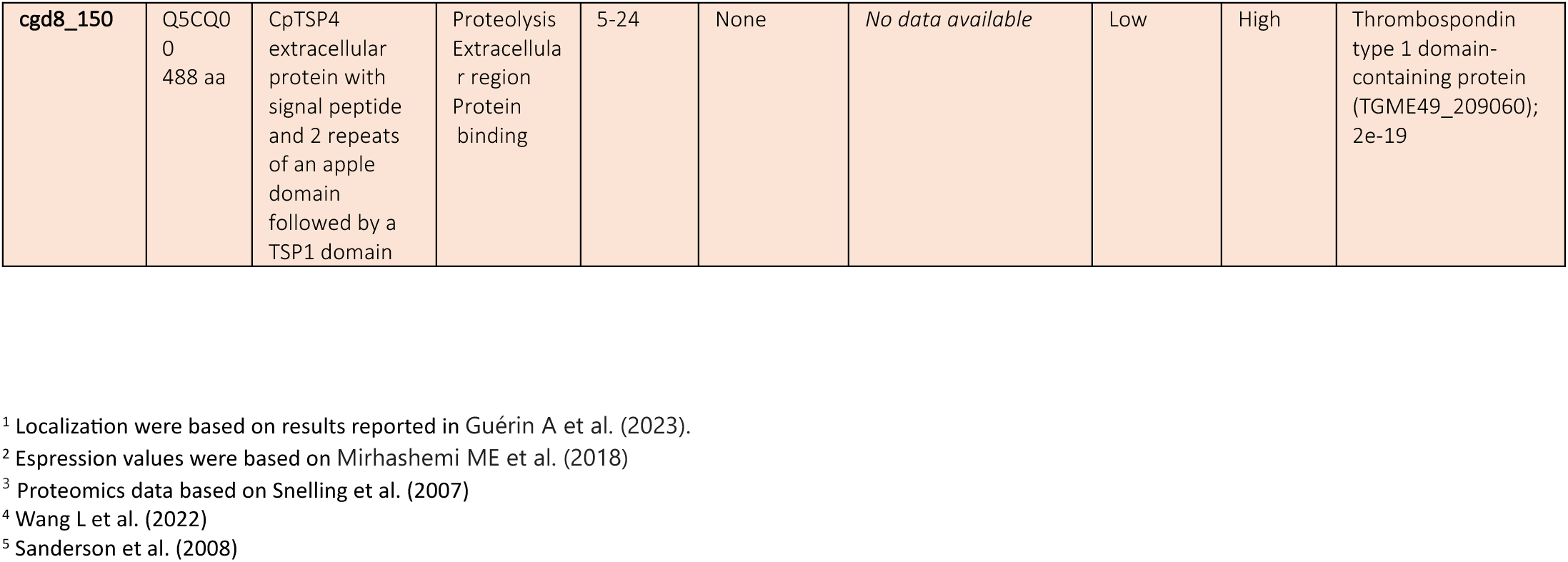
Putative targets of *C. parvum* rhomboids.

| CryptoDB ID | UniProt ID Length | Name (UniProt) | GO Terms | Position of TM (aa) | MS/MS data (Peptides in oocyst/sporozoite stage) | Localization in oocyst/sporozoite<sup>1</sup> | OOC/SPO<sup>2</sup> | Intracell. Stages<sup>2</sup> | Similarity in ToxoDB: Name and E value |
| --- | --- | --- | --- | --- | --- | --- | --- | --- | --- |
| Identified by similarity search and String analysis |  |  |  |  |  |  |  |  |  |
| **cgd2_1590** | Q5CTT8<br>488 aa | Extracellular protein with signal peptide, 5xEGF and apple domains | Proteolysis extracellular region<br>Calcium ion binding<br>Protein binding | 13-29 | None | Oocyst wall | High | Medium | Calcium-binding egf domain-containing protein CDBP2 (TGME49_318540); 4e-49 |
| **cgd6_670** | Q5CXK9<br>1607 aa | Extacellular protein with a signal peptide, clostripain like caspase/hemoglobinase domain, notch domain and 2 EGF domains | Integral component of membrane<br>Calcium ion binding | 1528-1550 | Yes (Insoluble excysted fraction<sup>3</sup>) | Oocyst wall | Low | Low | Calcium-binding egf domain-containing protein CDBP3 (TGME49_269930); 2e-62 |
| Identified only by similarity search |  |  |  |  |  |  |  |  |  |
| **cgd5_3420** | Q5CRC0<br>3869 aa | TRAP-C2 extracellular protein | Integral component of membrane | 3758-3780 | One (Oocyst wall<sup>4</sup>) | Inner membrane complex or Plasma membrane | Low | Low | Sushi domain (scr repeat) domain-containing protein (TGME49_223480) 1e-176 |
| **cgd2_3080** | Q5CTG7<br>391 aa | CpTSP10 protein | Integral component of membrane | 364-386 | None | Microneme | Low | High | Sushi domain (scr repeat) domain-containing protein (TGME49_223480); 6e-21 |
| **cgd5_4470** | Q5CQ18<br>656 aa | CpTSP7 extracellular membrane associated protein with a signal peptide followed by 2 TSP1 repeats, an EGF domain and a transmembrane region | Integral component of membrane | 563-582 | None | Microneme | Low | High | Sushi domain (scr repeat) domain-containing protein (TGME49_223480); 2e-35 |
| **cgd6_780** | Q5CXK1<br>625 aa | Putative extracellular protein | Integral component of membrane | Multispanning 17-36 599-621 | None | Microneme | High | Low | Sushi domain (scr repeat) domain-containing protein (TGME49_223480); 1e-24 |
| **cgd6_800** | Q5CXK0<br>457 aa | CpTSP9, extracellular protein with 3 TSP1 domains and an EGF domain | Integral component of membrane<br>calcium ion binding | 397-416 | None | Microneme | Low | High | Sushi domain (scr repeat) domain-containing protein (TGME49_223480); 1e-30 |
| **cgd1_3500** | Q5CSA5<br>687 aa | Thrombospondin related adhesive protein | Integral component of membrane | 622-644 | Many (Oocyst wall<sup>4</sup>) | PAM cl. 8 | Low | High | Thrombospondin type 1 domain-containing protein (TGME49_209060); 7e-32 |
| **cgd1_3510** | Q5CSA4<br>507 aa | TSP1 domain-containing protein TSP3 | *No data available* | 7-29 | Many (Oocyst wall<sup>4</sup>, Sporozoite<sup>5</sup>) | PAM cl. 8 | Low | High | Thrombospondin type 1 domain-containing protein (TGME49_209060); 2e-24 |
| **cgd8_150** | Q5CQ00<br>488 aa | CpTSP4 extracellular protein with signal peptide and 2 repeats of an apple domain followed by a TSP1 domain | Proteolysis Extracellular region Protein binding | 5-24 | None | *No data available* | Low | High | Thrombospondin type 1 domain-containing protein (TGME49_209060); 2e-19 |

<sup>1</sup> Localization were based on results reported in Guérin A et al. (2023).
<sup>2</sup> Expression values were based on Mirhashemi ME et al. (2018)
<sup>3</sup> Proteomics data based on Snelling et al. (2007)
<sup>4</sup> Wang L et al. (2022)
<sup>5</sup> Sanderson et al. (2008)

In parallel, we also explored the functional protein-protein interaction revealed by String to identify additional candidates for the rhomboid cleavage and the interaction maps generated by the String analysis were reported as supplementary material (Supplementary Figure S5). We listed 11 proteins as potential substrates considering only the proteins with a direct interaction with a rhomboid. It is to mention that 9 out of the 11 proteins of this list were attributed to interact with similar probability to each of the three *C. parvum* rhomboids.

Altogether, we listed 10 proteins as potential rhomboid substrates, with the following information (Table 2): gene ontology annotation which indicates the presumptive molecular function, location and/or involvement in a biological process. When available, we also included the MS/MS proteomic data and the localization in sporozoites by hiperLOPIT (Guérin et al., 2023). The approximate expression level was estimated based on mRNA in oocyst/sporozoite and intracellular stages (Mirhashemi et al., 2018).

All these proteins showed recognizable domains that indicated a putative function, but it was also evident that these *C. parvum* proteins could be phylogenetically classified in three orthologous groups based on the high similarity with specific proteins in *T. gondii* (last column on the right in Table 2).

Two of these proteins, namely cgd2_1590 and cgd6_670, share a calcium-binding EGF domain and the similarity with two related but different *T. gondii* proteins, namely CBDP2 and CBD3 (Wang et al., 2023). Interestingly, these two proteins were independently identified by similarity and String analysis.

The larger phylogenetic group, 5 proteins in total, was instead related to the Sushi domain-containing protein (TGME49_223480). This domain, also named CCP for complement control protein or SCR for short consensus repeats, has a role of complement inhibitor (Gialeli et al., 2018). Importantly, 4 of these 5 proteins have been localized in micronemes of sporozoites, thereby they sharing their localization with CpRom3 (Gao et al., 2021).

Finally, a small group of three proteins was highly related to a member of a TRAP protein family, namely the thrombospondin type 1 domain-containing protein (TGME49_209060). Furthermore, two of them, precisely cgd1_3500 and cgd1_3510, have been mapped (Guérin et al., 2023) in the same spatial cluster (PAM cl. 8) and identified by multiple peptides in proteomics experiments both in the oocyst wall (Wang et al., 2022) and in sporozoites before the excystation (Sanderson et al., 2008) but not in excysted sporozoites as expected for early cleaved and secreted proteins. Therefore, these two TRAP proteins represented good candidates as precociously exposed adhesins to contact the receptors on the host cell surface.

## Discussion

This is the first comprehensive study on *C. parvum* rhomboids, and it is the first effort to compare these proteins in terms of phylogenesis, structural characteristics and cell localization with the other apicomplexan rhomboids. Moreover, this study also identifies the putative substrates of these membrane proteases. In *C. parvum*, three distinct rhomboids are co-expressed in the sporozoite stage but each with its own specific sub-cellular distribution. Two of them, CpRom1 and CpRom3, apparently disappear after the formation of PV, whereas CpRom2, persist in the PV after the host cell invasion. The differences in the localization as well as the persistence or not in the intracellular stages indicate a different role in the parasite Biology.

The genes for CpRom1 and CpRom2 are constituted of single exons whereas the gene encoding for CpRom2 is composed of two exons. Genes composed of a single exon are the most common in *C. parvum*, since this species is characterized by a very compact genome (Abrahamsen et al., 2004). At the protein level these three rhomboids have significant differences in size, even if the genetic similarities show that each of the three *C. parvum* rhomboids derives from a limited number of ancestral paralogues, probably two, that have branched out in the evolution of Apicomplexa (Figure 2). Remarkably, *C. parvum* as well as other *Cryptosporidium* species lacks a mitochondrial PARL-like rhomboid.

The alignments of *C. parvum* rhomboids with functional rhomboids of *P. falciparum* and *T. gondii* reveal the presence of two distinguishable clusters that can be classified based on a short aminoacidic consensus around the catalytic dyad. The first cluster includes CpRom1 and CpRom2 and comprises big rhomboids larger than 400 aa and, among them, CpRom1 is the largest rhomboid ever described (990 aa). This cluster shows a conserved triad GSS at the fourth TM and an AHXGG consensus at the sixth TM. Functionally, PfRom4, TgRom4 and TgRom5, which belong to this group, are involved in adhesins’ cleavage suggesting a similar role for CpRom1 and CpRom2.

Differently, CpRom3 is related to a second cluster that includes PfRom1, PfRom3, TgRom1, TgRom2 and TgRom3. This group is characterized by a quadruple GAST aminoacidic sequence at the fourth TM and the GHIGG consensus at the sixth TM and it is constituted by smaller proteins of a maximum of 300 aa in length. Also PfRom1, which is included in this cluster, has a demonstrated capacity to cleave adhesins (Baker et al., 2006) and probably plays a role in the erythrocyte invasion (Singh et al., 2007). Differently, the roles of TgRom1, TgRom2 and TgRom3 are still uncertain even if their expression profiles are differentiated during the *T. gondii* life cycle (Brossier et al., 2005). Four rhomboids of this group, among them PfRom1 and CpRom3, also share a short consensus (FF or FPHF) that allows the Golgi sorting of these proteins to the destination organelle (i. e. the micronemes for CpRom3).

Based on the alignments and the TM domains, we have sketched the three *C. parvum* rhomboids and the PARL-related CmRom4 of *C. muris* for comparison (Figure 3). In this scheme, CpRom1 is presented with an additional 7^th^ TM domain and in this updated version the long N-terminus remains exposed on the external surface of the membrane as previously demonstrated (Bertuccini et al., 2024). Overall, all *C. parvum* rhomboids can be classified as mixed-secretase with 7 TM domain, a group in which functional apicomplexan rhomboids except for PARLs are included (Lemberg and Freeman, 2007).

As mentioned above, the similarity searches in *Cryptosporidium* spp reveal that *C. parvum* lacks a PARL-like gene, although this subclass of rhomboids is instead present in *C. muris* and *C. andersoni.* It is worth mentioning that *C. muris*, besides a PARL-like rhomboid, also conserves a functional mitochondrion (Uni et al., 1987). In addition, both *C. muris* and *C. andersoni* affect the stomach of their hosts and are thus indicated as “gastric” species. Differently, *C. parvum* and all the other species used for the genome comparison (Figure 4B) are all parasites of the small intestine and can be considered as “intestinal” species. The loss of the single mitochondrion has been a distinctive step in the evolution of this genus since it establishes a remarkable difference among the *Cryptosporidium* species and even more with the other apicomplexans. Indeed, *C. parvum* and other intestinal species lack an aerobic metabolism, whereas this remains in *C. muris* (Uni et al., 1987) and *C. andersoni* (Liu et al., 2016). These observations raise questions about the role of aerobic metabolism and the mitochondrion in the gastric *Cryptosporidium* species. Could the mitochondrion as well as the aerobic metabolism simply be a residual inheritance from the ancestors of this taxonomic group? In line with this view, genetic evidence supports such hypothesis since *C. muris* is more closely related to the *Cryptosporidium* ancestor (Xiao et al., 1999). Alternatively, is it possible that gastric species have retained this metabolism to survive in the stomach? Regardless of these questions, the precise excision of PARL-like genes in intestinal *Cryptosporidium* species without the modification of the chromosomal syntenic structure is worthy of note.

The expression of the recombinant forms of *C. parvum* rhomboids in *E. coli* has been feasible in denaturing conditions and the preparation of native proteins included in a cell membrane has been possible for 6h-CpRom2 and 6h-CpRom3 by means the expression in dedicated *E. coli* strains. Differently, the recombinant 6h-CpRom1 was always accumulated in insoluble inclusion bodies regardless of the *E. coli* strains (Supplementary Figure S1). It is reasonable that the presence of multi-spanning hydrophobic TM domains, combined with the lack of a specific transit route to the bacterial membrane, makes it difficult to produce well-conformed *C. parvum* rhomboids in bacteria. Indeed, the fractionations of the lysates from the recombinant strains have demonstrated the prevalence of insoluble forms of these proteins in bacterial cells. This experiment has also shown that the allocation in the biological membrane is required to obtain integer forms of 6h-CpRom2 and 6h-CpRom3. Nevertheless, the purification of recombinant peptides derived from these rhomboids has been possible in denaturing conditions thus allowing the production of specific mouse antisera.

CpRom2 and CpRom3 are expressed in intact oocysts and in excysted sporozoites similarly to CpRom1 (Bertuccini et al., 2024). Hence the three *C. parvum* rhomboids are co-present at least in the early stage of the life cycle of the parasite. In the oocysts/sporozoite stage, CpRom2 has been identified as a unique band of approximately 70 kDa, and the difference from the expected molecular weight of 51 kDa may be due to post-translational modifications, such as glycosylation (Figure 6A). Indeed, the recombinant 6h-CpRom2 synthetized in *E. coli* and localised in the membrane fraction is approximately of 50 kDa according to the expected molecular weight (Supplementary Figure S1). The amount of this protein remains unchanged from the oocyst to the excysted sporozoite, up to one hour after the excystation starts (Figure 7A). Differently, CpRom3 shown three different forms that vary in quantity during the excystation process (Figure 7B). The three forms are represented by a larger band of approximately 65-70 kDa in size, an intermediate band of approximately 35 kDa and a smaller one of roughly 30 kDa (Figure 7B). The 35 kDa band, which agrees with the expected molecular weight, appears only 30 minutes after the excystation and increases up to one-hour post-excystation. In contrast, the 30 kDa band that appears abundant in the oocysts and decreases in the excysted sporozoites. However, the largest 70 kDa band is continuously present from the oocysts stage to the excysted sporozoites after 1 hour. It is plausible that the largest 70 kDa band, which is approximately twice the expected molecular weight, may result from a paradoxical behaviour of certain rhomboids that can assume a dimeric or even multimeric forms in presence of detergents (i. e. SDS) (Sampathkumar et al., 2012). Indeed, the recombinant 6h-CpRom3 showed a prevalent band of 35 kDa, accordingly with the expected molecular weight, when localised in the *E. coli* cell membrane (Supplementary Figure S1). Hypothetically, the largest band (70 kDa) and the smallest band observed in the inactive oocysts (30kDa) could be the residual CpRom3 accumulated during the previous parasitic stages, whereas the 35 kDa band might represent the newly synthesized protein after the excystation. Overall, CpRom3 undergoes dynamic transformations at least during the early stage of the life cycle that modify forms and amounts of this protein.

We have then explored the localization of *C. parvum* rhomboids in the excysted sporozoites and showed that these proteins have a different spatial distribution.

Indeed, CpRom1 has a dual allocation: anterior to the nucleus and separately at the posterior end of the parasite. In the apical pole, the red labelling of this protein appears internal to the apical complex and not on the surface of the parasite, and the colocalization with the sporozoite antigens antiserum (Trasarti et al., 2007) that brightly labels in green the surface of sporozoites supports this interpretation (Figure 8C-8D). Notably, this CpRom1 distribution is consistent with a reported localization on the inner membrane complex (IMC), still determined with a different method (Guérin et al., 2023). On the other hand, anti-CpRom1 antibodies also intensely stain in red a thickening at the posterior pole of the sporozoites (Figure 8D). This dual localization has been confirmed by immunolocalization at the ultrastructural level with electron microscopy, as the gold particles have been observed inside the apical complex in the area around micronemes as well as at the posterior pole of sporozoites (Figure 9). A retro-grade route taken by CpRom1 from the apical complex towards the posterior pole of the parasite would explain this peculiar distribution. Actually, the CpRom1 distribution recalls that of TgRom5, which is the rhomboid more closely related to CpRom1. Indeed, TgRom5 converges towards the posterior pole to remove adhesins that would interfere with the parasite ingress in the host cell (Brossier et al., 2005).

Differently, CpRom2 is exclusively distributed around the apical complex (Figures 9-10) and the green labelling of this protein has always been observed anterior to the nucleus. Moreover, some of the not-permeabilised sporozoites show a brilliant green fluorescence on the apical surface meaning that CpRom2, at least in part, is transferred to the apical surface of sporozoites early after the excystation.

The immunolocalization of *C. parvum* rhomboids in the intracellular stages in HCT8 cells has been limited to the first 48 hours of infection; however this experiment demonstrates that CpRom2 persists in the PV of the infected cells after the entry of the parasite. Hence, CpRom2 can be observed at intracellular stages of the parasite (Figure 11). This suggests that, differently from CpRom1 and CpRom3, CpRom2 plays a role in the early internal stages of the life cycle of the parasite.

The localization of CpRom3 is further different from that of CpRom1 and CpRom2, in fact this protein is distributed throughout the surface of the sporozoite from the anterior pole up to the posterior pole (Figure 11). The specific labelling also shows two points of thickening right on the extremities of the sporozoite (Figure 11E). This distribution is perfectly in accordance with previous subcellular localization that assigned this protein in the apical micronemes and on the pellicle of sporozoites (Gao et al., 2021). Overall, the localization in the micronemes and on the external membrane, as well as the high similarity with Pfrom1 (Baker et al., 2006), suggests that this rhomboid is probably involved in the adhesin cleavage along the entire sporozoite membrane.

The genetic manipulation of *C. parvum* to demonstrate functional interaction between a specific rhomboid and putative adhesins is still difficult; so the search for similarities with other apicomplexan adhesins is a feasible path to identify the putative targets of *C. parvum* rhomboids. The in-silico analysis to identify plausible substrates for the *C. parvum* rhomboids identified various proteins with adhesive domains that most probably are involved in the first phases of the progression toward and the adhesion to the host enterocytes. Some of these proteins share domains and high phylogenetic relations with well-known adhesive proteins just described in other Apicomplexa. In this context, it is to mention that the *C. parvum* genome did not reveal any protein related to the AMA proteins, which are instead present in *P. falciparum* and in *T.gondii* (data not shown) and that are substrates of rhomboids in these parasites (Sheiner et al., 2010). On the other hand, all the identified *C. parvum* proteins are strictly related to orthologs in the *T.gondii* genome. Indeed, the two proteins with a calcium binding EGF domain also identified by String, namely cgd2_1590 and cgd6_670, are strictly related to CBDP2 and to CBDP3, which are two proteins of *T. gondii* with calcium-binding epidermal growth factor (EGF) domains. However, the role of these calcium-binding epidermal growth factor (EGF) domain-containing proteins such as CBDP2 and to CBDP3 in *T. gondii* infection is still elusive (Wang et al., 2023).

Differently, the *C. parvum* proteins with a CCP domain may have an important function before the penetration of the sporozoite in the host cell. This search has identified a group of 5 proteins highly related to the Sushi domain (scr repeat) domain-containing protein (TGME49_223480) of *T. gondii* and 4 of these proteins have been localized in micronemes (Guérin A et al., 2023). Sushi domains are characteristic of mammalian proteins in the extracellular matrix and play a role in the complement regulation (Gialeli et al., 2018). Among the parasites, proteins with the Sushi domain are present not only in Apicomplexa but Sushi-provided proteins are also expressed in tissue-larvae of nematodes (Hewitson et al., 2013). It is plausible that these parasitic proteins may play a role in reducing the action of the complement factors against the parasites.

Finally, a smaller group of three proteins is represented by *C. parvum* proteins related to the thrombospondin type 1 domain-containing protein (TGME49_209060) of *T. gondii*. This protein belongs to the TRAP family and this type of multi-modular proteins is known to be necessary for the invasion of the host cell (Morahan et al., 2009). TRAP proteins have multiple adhesive domains that allow the adhesion to and the gliding over the host-cells (Paoletta and Wilkowsky, 2022). Two of these thrombospondin-related proteins, precisely cgd1_3500 and cgd1_3510, have been identified by multiple peptides in proteomics experiments both in the oocyst wall (Wang et al., 2022) and in sporozoites before the excystation (Sanderson et al., 2008), but not in excysted sporozoites (Snelling et al., 2007). Therefore, these adhesive proteins may represent precocious molecules exposed by the parasite to contact the receptors on the host cell surface. The identification of the *C. parvum* membrane proteins cleaved by rhomboids should be supported by dedicated proteomics experiments on the proteins secreted by sporozoites after the excystation and prior to the host cell invasion.

To conclude, this is the first comprehensive study on the three *C. parvum* rhomboids, and we have shown that these proteins are highly related to rhomboids of other Apicomplexa. It has been demonstrated that these proteases, specifically in *P. falciparum* and *T. gondii*, play a crucial role in removing adhesive proteins from the cell surface, favouring a proper positioning and entry before and during the host cell invasion (Brossier et al., 2005; Howell et al., 2005; Buguliskis et al., 2010). In fact, genetically modified parasites in which some rhomboid functions were abolished have a strongly reduced infectious capacity (Shen et al., 2014). We have also identified a decade of protein candidates as rhomboid substrates, and these proteins are functionally related to *T. gondii* and *P. falciparum* proteins involved in the immunoevasion or in the attack of the host cells. Overall, this study identifies a group of sporozoite proteins, namely rhomboids and their presumptive substrates, that can be targeted by a therapy such as specific antibodies to block the parasite entry in intestinal enterocytes.

## Acknowledgements

We are thankful to VEuPathDB for making available all the genomic, transcriptomic and proteomic data used in this study. We thank James U. Bowie for the gift of the *E. coli* EXP strains.

## References

1. Abrahamsen MS, Templeton TJ, Enomoto S, Abrahante JE, Zhu G, Lancto CA, Deng M, Liu C, Widmer G, Tzipori S, Buck GA, Xu P, Bankier AT, Dear PH, Konfortov BA, Spriggs HF, Iyer L, Anantharaman V, Aravind L, Kapur V. Complete genome sequence of the apicomplexan, Cryptosporidium parvum. Science. 2004 Apr 16;304(5669):441-5. doi: 10.1126/science.1094786. Epub 2004 Mar 25. PMID: 15044751.

2. Baker RP, Wijetilaka R, Urban S. Two Plasmodium rhomboid proteases preferentially cleave different adhesins implicated in all invasive stages of malaria. PLoS Pathog. 2006 Oct;2(10):e113. doi: 10.1371/journal.ppat.0020113. PMID: 17040128; PMCID: PMC1599764.

3. Baker RP, Young K, Feng L, Shi Y, Urban S. Enzymatic analysis of a rhomboid intramembrane protease implicates transmembrane helix 5 as the lateral substrate gate. Proc Natl Acad Sci U S A. 2007 May 15;104(20):8257–62. doi: 10.1073/pnas.0700814104. Epub 2007 Apr 26. PMID: 17463085; PMCID: PMC1895938.

4. Balendran T, Iddawela D, Lenadora S. Cryptosporidiosis in a Zoonotic Gastrointestinal Disorder Perspective: Present Status, Risk Factors, Pathophysiology, and Treatment, Particularly in Immunocompromised Patients. J Trop Med. 2024 Nov 5;2024:6439375. doi: 10.1155/2024/6439375. PMID: 39534184; PMCID: PMC11557182.

5. Bergbold N, Lemberg MK. Emerging role of rhomboid family proteins in mammalian biology and disease. Biochim Biophys Acta. 2013 Dec;1828(12):2840–8. doi: 10.1016/j.bbamem.2013.03.025. Epub 2013 Apr 3. PMID: 23562403.

6. Bertuccini L, Boussadia Z, Salzano AM, Vanni I, Passerò I, Nocita E, Scaloni A, Sanchez M, Sargiacomo M, Fiani ML, Tosini F. Unveiling *Cryptosporidium parvum* sporozoite-derived extracellular vesicles: profiling, origin, and protein composition. Front Cell Infect Microbiol. 2024 Apr 10;14:1367359. doi: 10.3389/fcimb.2024.1367359. PMID: 38660488; PMCID: PMC11039866.

7. Brossier F, Jewett TJ, Sibley LD, Urban S. A spatially localized rhomboid protease cleaves cell surface adhesins essential for invasion by Toxoplasma. Proc Natl Acad Sci U S A. 2005 Mar 15;102(11):4146–51. doi: 10.1073/pnas.0407918102. Epub 2005 Mar 7. PMID: 15753289; PMCID: PMC554800.

8. Buguliskis JS, Brossier F, Shuman J, Sibley LD. Rhomboid 4 (ROM4) affects the processing of surface adhesins and facilitates host cell invasion by Toxoplasma gondii. PLoS Pathog. 2010 Apr 22;6(4):e1000858. doi: 10.1371/journal.ppat.1000858. PMID: 20421941; PMCID: PMC2858701.

9. Carruthers VB. Proteolysis and Toxoplasma invasion. Int J Parasitol. 2006 May 1;36(5):595-600. doi: 10.1016/j.ijpara.2006.02.008. Epub 2006 Mar 13. PMID: 16600244.

10. Castellanos-Gonzalez A, Martinez-Traverso G, Fishbeck K, Nava S, White AC Jr. Systematic gene silencing identified Cryptosporidium nucleoside diphosphate kinase and other molecules as targets for suppression of parasite proliferation in human intestinal cells. Sci Rep. 2019 Aug 21;9(1):12153. doi: 10.1038/s41598-019-48544-z. PMID: 31434931; PMCID: PMC6704102.

11. Dulloo I, Muliyil S, Freeman M. The molecular, cellular and pathophysiological roles of iRhom pseudoproteases. Open Biol. 2019 Mar 29;9(3):190003. doi: 10.1098/rsob.190003. PMID: 30890028; PMCID: PMC6451368.

12. Fleming J, Magana P, Nair S, Tsenkov M, Bertoni D, Pidruchna I, Lima Afonso MQ, Midlik A, Paramval U, Žídek A, Laydon A, Kovalevskiy O, Pan J, Cheng J, Avsec Ž, BycroÇ C, Wong LH, Last M, Mirdita M, Steinegger M, Kohli P, Váradi M, Velankar S. AlphaFold Protein Structure Database and 3D-Beacons: New Data and Capabilities. J Mol Biol. 2025 Aug 1;437(15):168967. doi: 10.1016/j.jmb.2025.168967. Epub 2025 Jan 29. PMID: 40133787.

13. Gallagher JR, Matthews KA, Prigge ST. Plasmodium falciparum apicoplast transit peptides are unstructured in vitro and during apicoplast import. Traffic. 2011 Sep;12(9):1124–38. doi: 10.1111/j.1600-0854.2011.01232.x. Epub 2011 Jul 7. PMID: 21668595; PMCID: PMC3629917.

14. Gao X, Yin J, Wang D, Li X, Zhang Y, Wang C, Zhang Y, Zhu G. Discovery of New Microneme Proteins in *Cryptosporidium parvum* and Implication of the Roles of a Rhomboid Membrane Protein (CpROM1) in Host-Parasite Interaction. Front Vet Sci. 2021 Dec 13;8:778560. doi: 10.3389/fvets.2021.778560. PMID: 34966810; PMCID: PMC8710574.

15. Gialeli C, Gungor B, Blom AM. Novel potential inhibitors of complement system and their roles in complement regulation and beyond. Mol Immunol. 2018 Oct;102:73–83. doi: 10.1016/j.molimm.2018.05.023. Epub 2018 Jun 7. PMID: 30217334.

16. Guérin A, Strelau KM, Barylyuk K, Wallbank BA, Berry L, Crook OM, Lilley KS, Waller RF, Striepen B. Cryptosporidium uses multiple distinct secretory organelles to interact with and modify its host cell. Cell Host Microbe. 2023 Apr 12;31(4):650–664.e6. doi: 10.1016/j.chom.2023.03.001. Epub 2023 Mar 22. PMID: 36958336.

17. Hallgren J, Tsirigos DK, Pedersen MD, Almagro Armenteros JJA, Marcatili P, Nielsen H, Krogh A, Winther O. DeepTMHMM predicts alpha and beta transmembrane proteins using deep neural networks. (2022) https://doi.org/10.1101/2022.04.08.487609

18. Helmy YA, Hafez HM. Cryptosporidiosis: From Prevention to Treatment, a Narrative Review. Microorganisms. 2022 Dec 13;10(12):2456. doi: 10.3390/microorganisms10122456. PMID: 36557709; PMCID: PMC9782356.

19. Hewitson JP, Ivens AC, Harcus Y, Filbey KJ, McSorley HJ, Murray J, Bridgett S, Ashford D, Dowle AA, Maizels RM. Secretion of protective antigens by tissue-stage nematode larvae revealed by proteomic analysis and vaccination-induced sterile immunity. PLoS Pathog. 2013 Aug;9(8):e1003492. doi: 10.1371/journal.ppat.1003492. Epub 2013 Aug 15. PMID: 23966853; PMCID: PMC3744408.

20. Howell SA, Hackett F, Jongco AM, Withers-Martinez C, Kim K, Carruthers VB, Blackman MJ. Distinct mechanisms govern proteolytic shedding of a key invasion protein in apicomplexan pathogens. Mol Microbiol. 2005 Sep;57(5):1342–56. doi: 10.1111/j.1365-2958.2005.04772.x. PMID: 16102004.

21. Jumper J, Evans R, Pritzel A, Green T, Figurnov M, Ronneberger O, Tunyasuvunakool K, Bates R, Žídek A, Potapenko A, Bridgland A, Meyer C, Kohl SAA, Ballard AJ, Cowie A, Romera-Paredes B, Nikolov S, Jain R, Adler J, Back T, Petersen S, Reiman D, Clancy E, Zielinski M, Steinegger M, Pacholska M, Berghammer T, Bodenstein S, Silver D, Vinyals O, Senior AW, Kavukcuoglu K, Kohli P, Hassabis D. Highly accurate protein structure prediction with AlphaFold. Nature. 2021 Aug;596(7873):583-589. doi: 10.1038/s41586-021-03819-2. Epub 2021 Jul 15. PMID: 34265844; PMCID: PMC8371605.

22. Khalil IA, Troeger C, Rao PC, Blacker BF, Brown A, Brewer TG, Colombara DV, De Hostos EL, Engmann C, Guerrant RL, Haque R, Houpt ER, Kang G, Korpe PS, Kotloff KL, Lima AAM, Petri WA Jr, Platts-Mills JA, Shoultz DA, Forouzanfar MH, Hay SI, Reiner RC Jr, Mokdad AH. Morbidity, mortality, and long-term consequences associated with diarrhoea from Cryptosporidium infection in children younger than 5 years: a meta-analyses study. Lancet Glob Health. 2018 Jul;6(7):e758–e768. doi: 10.1016/S2214-109X(18)30283-3. PMID: 29903377; PMCID: PMC6005120.

23. Kreutzberger AJB, Urban S. Single-Molecule Analyses Reveal Rhomboid Proteins Are Strict and Functional Monomers in the Membrane. Biophys J. 2018 Nov 6;115(9):1755–1761. doi: 10.1016/j.bpj.2018.09.024. Epub 2018 Oct 2. PMID: 30342748; PMCID: PMC6224778.

24. Kumar S, Stecher G, Suleski M, Sanderford M, Sharma S, Tamura K. MEGA12: Molecular Evolutionary Genetic Analysis Version 12 for Adaptive and Green Computing. Mol Biol Evol. 2024 Dec 6;41(12):msae263. doi: 10.1093/molbev/msae263. PMID: 39708372; PMCID: PMC11683415.

25. Lastun VL, Grieve AG, Freeman M. Substrates and physiological functions of secretase rhomboid proteases. Semin Cell Dev Biol. 2016 Dec;60:10–18. doi: 10.1016/j.semcdb.2016.07.033. Epub 2016 Aug 4. PMID: 27497690.

26. Lemberg MK, Freeman M. Functional and evolutionary implications of enhanced genomic analysis of rhomboid intramembrane proteases. Genome Res. 2007 Nov;17(11):1634–46. doi: 10.1101/gr.6425307. Epub 2007 Oct 15. PMID: 17938163; PMCID: PMC2045146.

27. Lemberg MK, Strisovsky K. Maintenance of organellar protein homeostasis by ER-associated degradation and related mechanisms. Mol Cell. 2021 Jun 17;81(12):2507–2519. doi: 10.1016/j.molcel.2021.05.004. Epub 2021 Jun 8. PMID: 34107306.

28. Li M, Zhang X, Gong P, Li J. Cryptosporidium parvum rhomboid1 has an activity in microneme protein CpGP900 cleavage. Parasit Vectors. 2016 Aug 8;9(1):438. doi: 10.1186/s13071-016-1728-6. PMID: 27502595; PMCID: PMC4977710.

29. Liu S, Roellig DM, Guo Y, Li N, Frace MA, Tang K, Zhang L, Feng Y, Xiao L. Evolution of mitosome metabolism and invasion-related proteins in Cryptosporidium. BMC Genomics. 2016 Dec 8;17(1):1006. doi: 10.1186/s12864-016-3343-5. PMID: 27931183; PMCID: PMC5146892.

30. Lysyk L, Brassard R, Touret N, Lemieux MJ. PARL Protease: A Glimpse at Intramembrane Proteolysis in the Inner Mitochondrial Membrane. J Mol Biol. 2020 Aug 21;432(18):5052–5062. doi: 10.1016/j.jmb.2020.04.006. Epub 2020 Apr 19. PMID: 32320686.

31. Massey-Gendel E, Zhao A, Boulting G, Kim HY, Balamotis MA, Seligman LM, Nakamoto RK, Bowie JU. Genetic selection system for improving recombinant membrane protein expression in E. coli. Protein Sci. 2009 Feb;18(2):372–83. doi: 10.1002/pro.39. PMID: 19165721; PMCID: PMC2708063.

32. Mirhashemi ME, Noubary F, Chapman-Bonofiglio S, Tzipori S, Huggins GS, Widmer G. Transcriptome analysis of pig intestinal cell monolayers infected with Cryptosporidium parvum asexual stages. Parasit Vectors. 2018 Mar 12;11(1):176. doi: 10.1186/s13071-018-2754-3. PMID: 29530089; PMCID: PMC5848449.

33. Morahan BJ, Wang L, Coppel RL. No TRAP, no invasion. Trends Parasitol. 2009 Feb;25(2):77-84. doi: 10.1016/j.pt.2008.11.004. Epub 2008 Dec 26. PMID: 19101208.

34. Omasits U, Ahrens CH, Müller S, Wollscheid B. Protter: interactive protein feature visualization and integration with experimental proteomic data. Bioinformatics. 2014 Mar 15;30(6):884–6. doi: 10.1093/bioinformatics/bZ607. Epub 2013 Oct 24. PMID: 24162465.

35. Paoletta MS, Wilkowsky SE. Thrombospondin Related Anonymous Protein Superfamily in Vector-Borne Apicomplexans: The Parasite’s Toolkit for Cell Invasion. Front Cell Infect Microbiol. 2022 Apr 6;12:831592. doi: 10.3389/fcimb.2022.831592. PMID: 35463644; PMCID: PMC9019593.

36. Rugarabamu G, Marq JB, Guérin A, Lebrun M, Soldati-Favre D. Distinct contribution of Toxoplasma gondii rhomboid proteases 4 and 5 to micronemal protein protease 1 activity during invasion. Mol Microbiol. 2015 Jul;97(2):244–62. doi: 10.1111/mmi.13021. Epub 2015 May 9. PMID: 25846828.

37. Ryan U, Zahedi A, Feng Y, Xiao L. An Update on Zoonotic *Cryptosporidium* Species and Genotypes in Humans. Animals (Basel). 2021 Nov 19;11(11):3307. doi: 10.3390/ani11113307. PMID: 34828043; PMCID: PMC8614385.

38. Sambrook, J., Fritsch, E. F., & Maniatis, T. (1989). Molecular cloning: A laboratory manual (2nd ed.). Cold Spring Harbor Laboratory Press.

39. Sampathkumar P, Mak MW, Fischer-Witholt SJ, Guigard E, Kay CM, Lemieux MJ. Oligomeric state study of prokaryotic rhomboid proteases. Biochim Biophys Acta. 2012 Dec;1818(12):3090–7. doi: 10.1016/j.bbamem.2012.08.004. Epub 2012 Aug 18. PMID: 22921757.

40. Sampathkumar P, Mak MW, Fischer-Witholt SJ, Guigard E, Kay CM, Lemieux MJ. Oligomeric state study of prokaryotic rhomboid proteases. Biochim Biophys Acta. 2012 Dec;1818(12):3090–7. doi: 10.1016/j.bbamem.2012.08.004. Epub 2012 Aug 18. PMID: 22921757.

41. Sanderson SJ, Xia D, Prieto H, Yates J, Heiges M, Kissinger JC, Bromley E, Lal K, Sinden RE, Tomley F, Wastling JM. Determining the protein repertoire of Cryptosporidium parvum sporozoites. Proteomics. 2008 Apr;8(7):1398–414. doi: 10.1002/pmic.200700804. PMID: 18306179; PMCID: PMC2770187.

42. Sanger, F., Nicklen, S., & Coulson, A. R. (1977). DNA sequencing with chain-terminating inhibitors. Proceedings of the National Academy of Sciences, 74(12), 5463–5467.

43. Sheiner L, Dowse TJ, Soldati-Favre D. Identification of trafficking determinants for polytopic rhomboid proteases in Toxoplasma gondii. Traffic. 2008 May;9(5):665–77. doi: 10.1111/j.1600-0854.2008.00736.x. Epub 2008 Mar 10. PMID: 18346213.

44. Sheiner L, Santos JM, Klages N, Parussini F, Jemmely N, Friedrich N, Ward GE, Soldati-Favre D. Toxoplasma gondii transmembrane microneme proteins and their modular design. Mol Microbiol. 2010 Aug;77(4):912–29. doi: 10.1111/j.1365-2958.2010.07255.x. Epub 2010 Jun 9. PMID: 20545864; PMCID: PMC2982875.

45. Shen B, Buguliskis JS, Lee TD, Sibley LD. Functional analysis of rhomboid proteases during Toxoplasma invasion. mBio. 2014 Oct 21;5(5):e01795–14. doi: 10.1128/mBio.01795-14. PMID: 25336455; PMCID: PMC4212836.

46. Sibley LD. The roles of intramembrane proteases in protozoan parasites. Biochim Biophys Acta. 2013 Dec;1828(12):2908–15. doi: 10.1016/j.bbamem.2013.04.017. PMID: 24099008; PMCID: PMC3793208.

47. Singh S, Plassmeyer M, Gaur D, Miller LH. Mononeme: a new secretory organelle in Plasmodium falciparum merozoites identified by localization of rhomboid-1 protease. Proc Natl Acad Sci U S A. 2007 Dec 11;104(50):20043–8. doi: 10.1073/pnas.0709999104. Epub 2007 Nov 28. PMID: 18048320; PMCID: PMC2148419.

48. Snelling WJ, Lin Q, Moore JE, Millar BC, Tosini F, Pozio E, Dooley JS, Lowery CJ. Proteomics analysis and protein expression during sporozoite excystation of Cryptosporidium parvum (Coccidia, Apicomplexa). Mol Cell Proteomics. 2007 Feb;6(2):346–55. doi: 10.1074/mcp.M600372-MCP200. Epub 2006 Nov 23. PMID: 17124246.

49. Strisovsky K, Sharpe HJ, Freeman M. Sequence-specific intramembrane proteolysis: identification of a recognition motif in rhomboid substrates. Mol Cell. 2009 Dec 25;36(6):1048–59. doi: 10.1016/j.molcel.2009.11.006. PMID: 20064469; PMCID: PMC2941825.

50. Strisovsky K. Structural and mechanistic principles of intramembrane proteolysis--lessons from rhomboids. FEBS J. 2013 Apr;280(7):1579–603. doi: 10.1111/febs.12199. Epub 2013 Mar 20. PMID: 23432912.

51. Tichá A, Collis B, Strisovsky K. The Rhomboid Superfamily: Structural Mechanisms and Chemical Biology Opportunities. Trends Biochem Sci. 2018 Sep;43(9):726–739. doi: 10.1016/j.tibs.2018.06.009. Epub 2018 Jul 25. PMID: 30055896.

52. Tosini F, Agnoli A, Mele R, Gomez Morales MA, Pozio E. A new modular protein of Cryptosporidium parvum, with ricin B and LCCL domains, expressed in the sporozoite invasive stage. Mol Biochem Parasitol. 2004 Mar;134(1):137–47. doi: 10.1016/j.molbiopara.2003.11.014. PMID: 14747151.

53. Trasarti E, Pizzi E, Pozio E, Tosini F. The immunological selection of recombinant peptides from Cryptosporidium parvum reveals 14 proteins expressed at the sporozoite stage, 7 of which are conserved in other apicomplexa. Mol Biochem Parasitol. 2007 Apr;152(2):159-69. doi: 10.1016/j.molbiopara.2006.12.010. Epub 2007 Jan 7. PMID: 17267054.

54. Truong Q, Ferrari BC. Quantitative and qualitative comparisons of Cryptosporidium faecal purification procedures for the isolation of oocysts suitable for proteomic analysis. Int J Parasitol. 2006 Jun;36(7):811–9. doi: 10.1016/j.ijpara.2006.02.023. Epub 2006 Mar 31. PMID: 16696982.

55. Uni S, Iseki M, Maekawa T, Moriya K, Takada S. Ultrastructure of Cryptosporidium muris (strain RN 66) parasitizing the murine stomach. Parasitol Res. 1987;74(2):123–32. doi: 10.1007/BF00536023. PMID: 2964037.

56. Wang L, Wang Y, Cui Z, Li D, Li X, Zhang S, Zhang L. Enrichment and proteomic identification of Cryptosporidium parvum oocyst wall. Parasit Vectors. 2022 Sep 23;15(1):335. doi: 10.1186/s13071-022-05448-8. PMID: 36151578; PMCID: PMC9508764.

57. Wang XC, Li TT, Elsheikha HM, Zheng XN, Zhao DY, Wang JL, Wang M, Zhu XQ. Effect of deleting four Toxoplasma gondii calcium-binding EGF domain-containing proteins on parasite replication and virulence. Parasitol Res. 2023 Feb;122(2):441–450. doi: 10.1007/s00436-022-07739-6. Epub 2022 Dec 6. PMID: 36471092.

58. Xiao L, Escalante L, Yang C, Sulaiman I, Escalante AA, Montali RJ, Fayer R, Lal AA. Phylogenetic analysis of Cryptosporidium parasites based on the small-subunit rRNA gene locus. Appl Environ Microbiol. 1999 Apr;65(4):1578–83. doi: 10.1128/AEM.65.4.1578-1583.1999. PMID: 10103253; PMCID: PMC91223.

59. Zhou XW, Blackman MJ, Howell SA, Carruthers VB. Proteomic analysis of cleavage events reveals a dynamic two-step mechanism for proteolysis of a key parasite adhesive complex. Mol Cell Proteomics. 2004 Jun;3(6):565–76. doi: 10.1074/mcp.M300123-MCP200. Epub 2004 Feb 24. PMID: 14982962.

