## Supplementary figures and images for "Exploring the Repertoire of Rhomboid Proteases in *Cryptosporidium parvum* Parasite: Phylogenesis, Structural motifs and Cellular Localization in Sporozoite Cells"

### Supplementary Figure S1

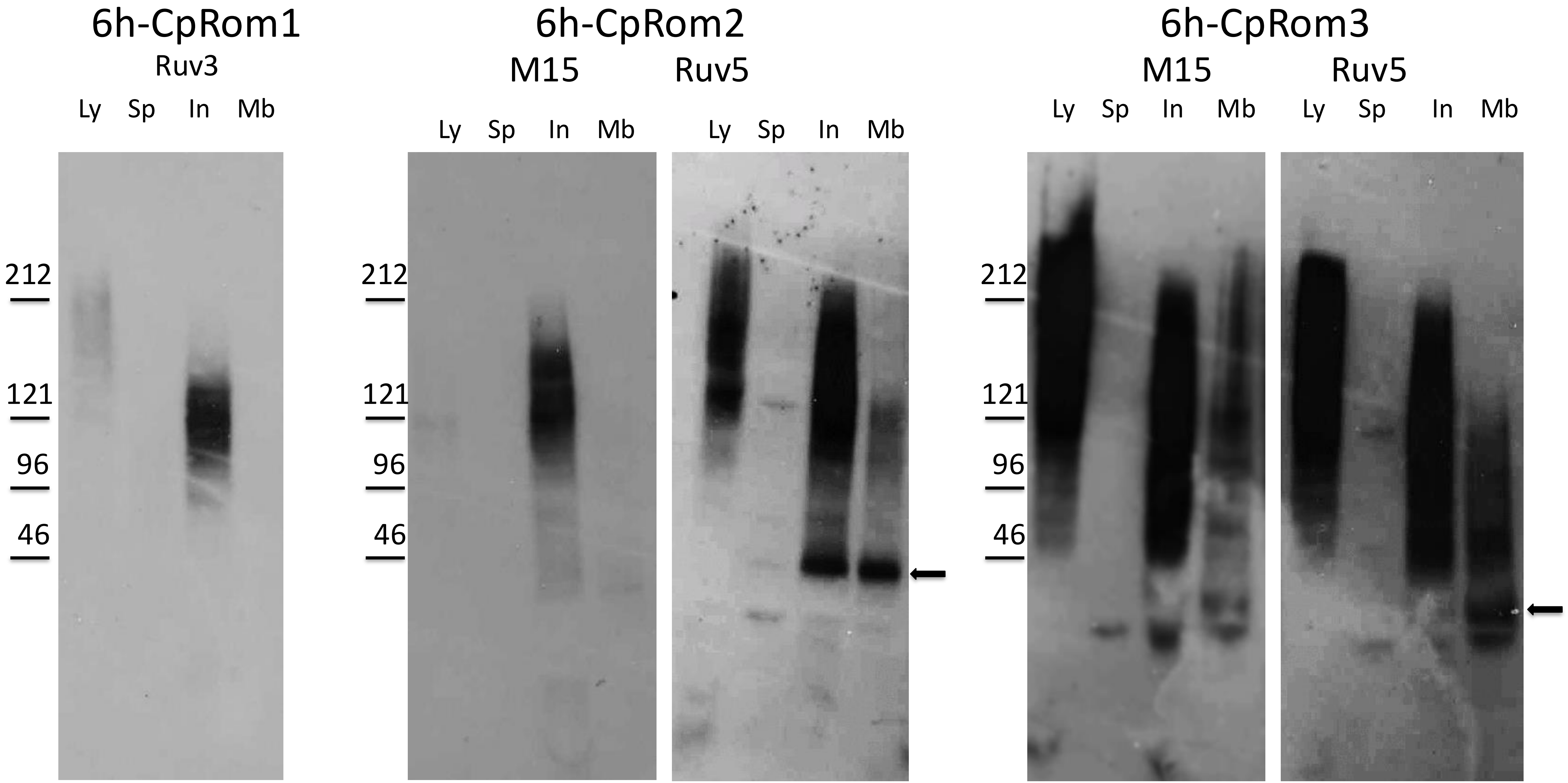

### Supplementary Figure S2

#### C. parvum rhomboid constructs in expression vector pQE30

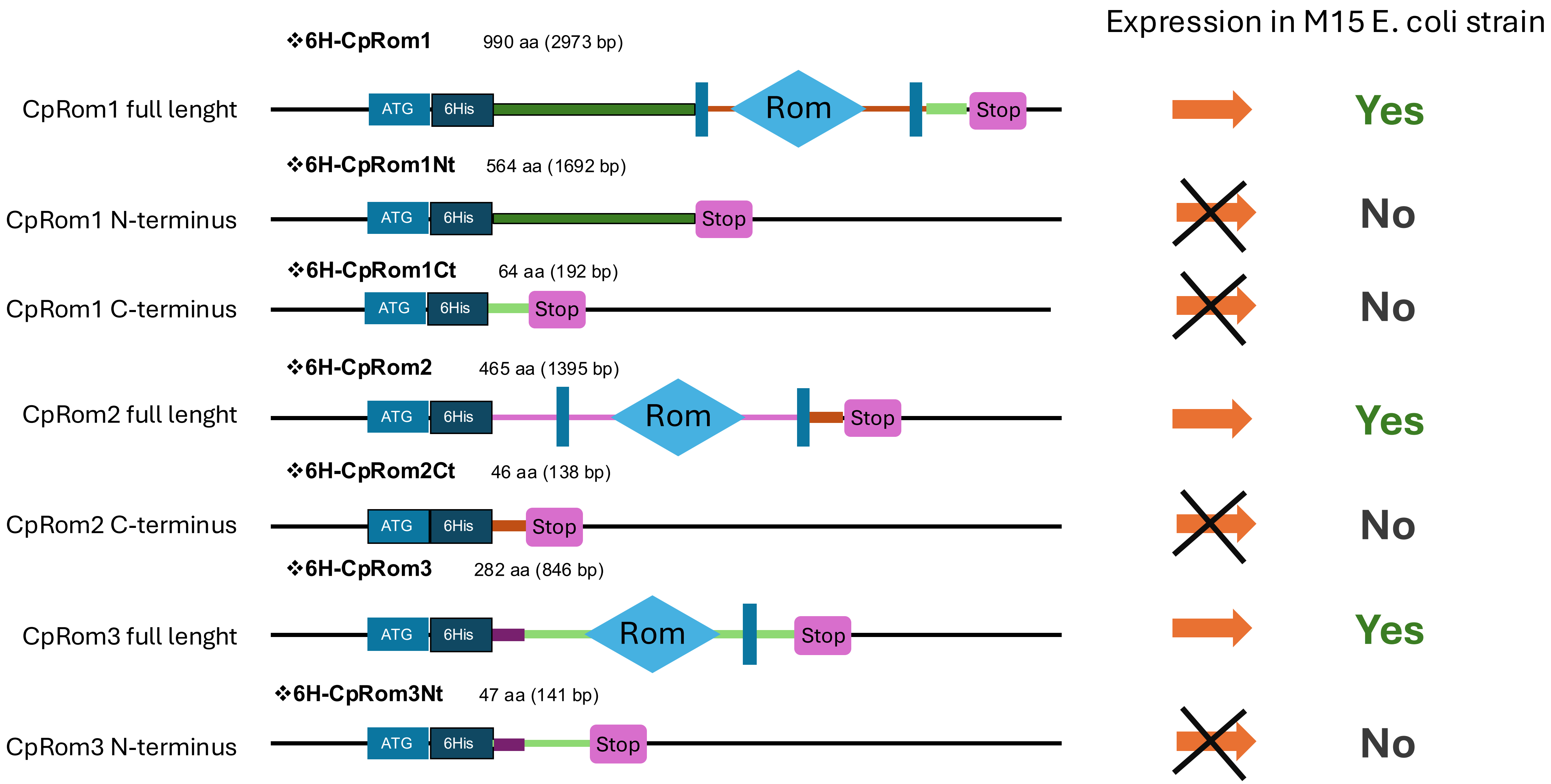

### Supplementary Figure S3

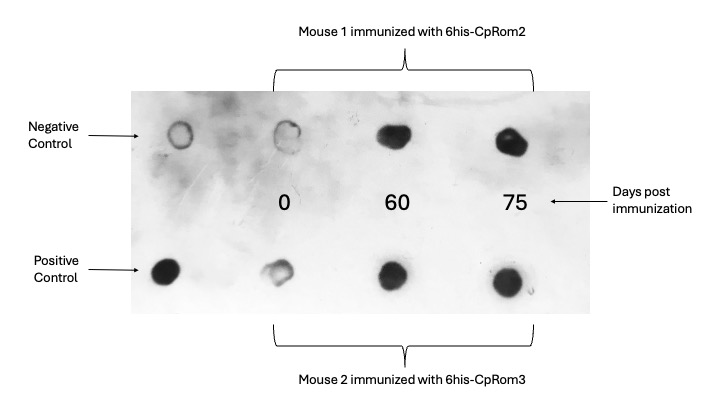

### Supplementary Figure S4

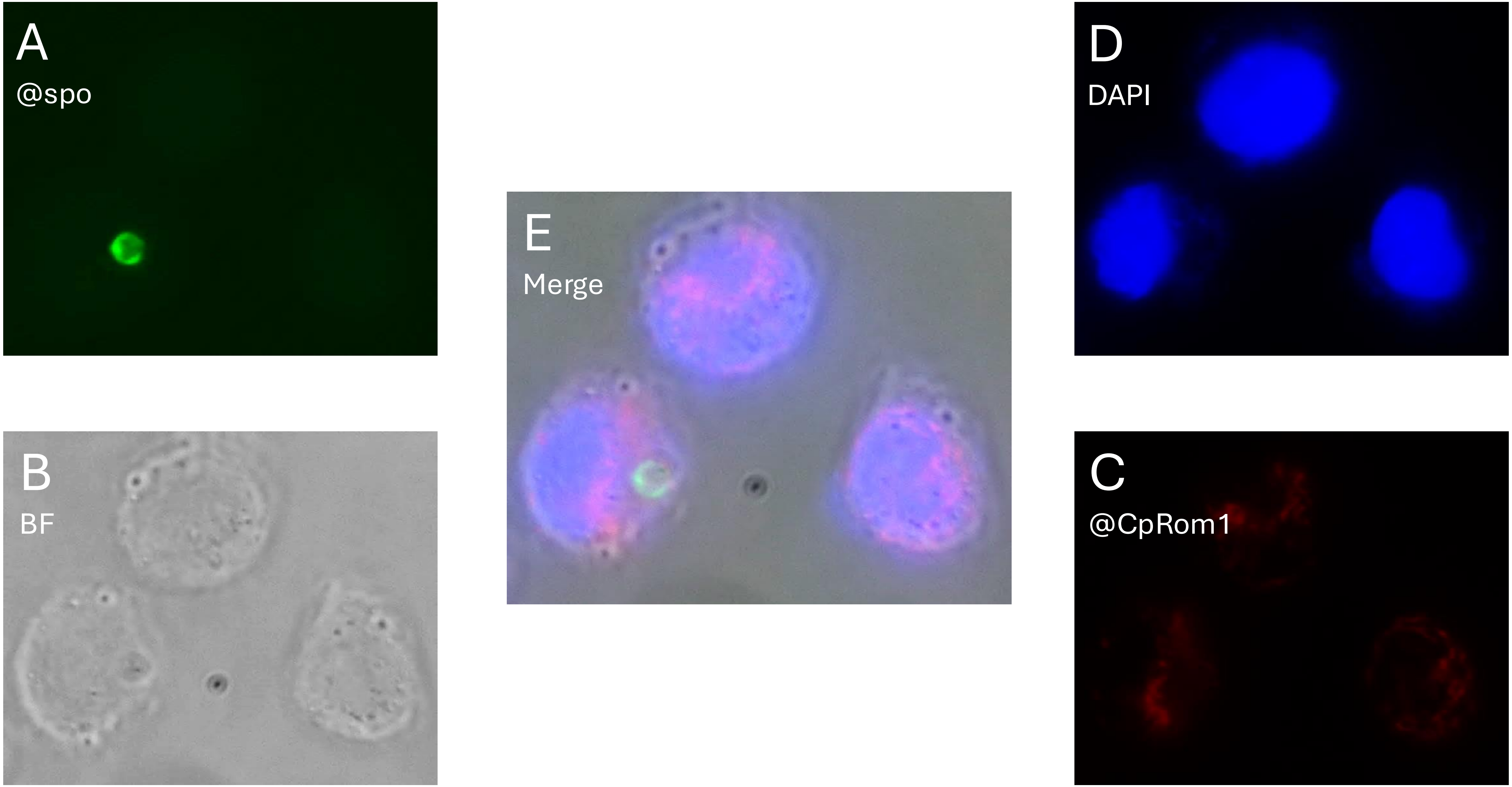
