## Supplementary Figure S5 for "Exploring the Repertoire of Rhomboid Proteases in *Cryptosporidium parvum* Parasite: Phylogenesis, Structural motifs and Cellular Localization in Sporozoite Cells"

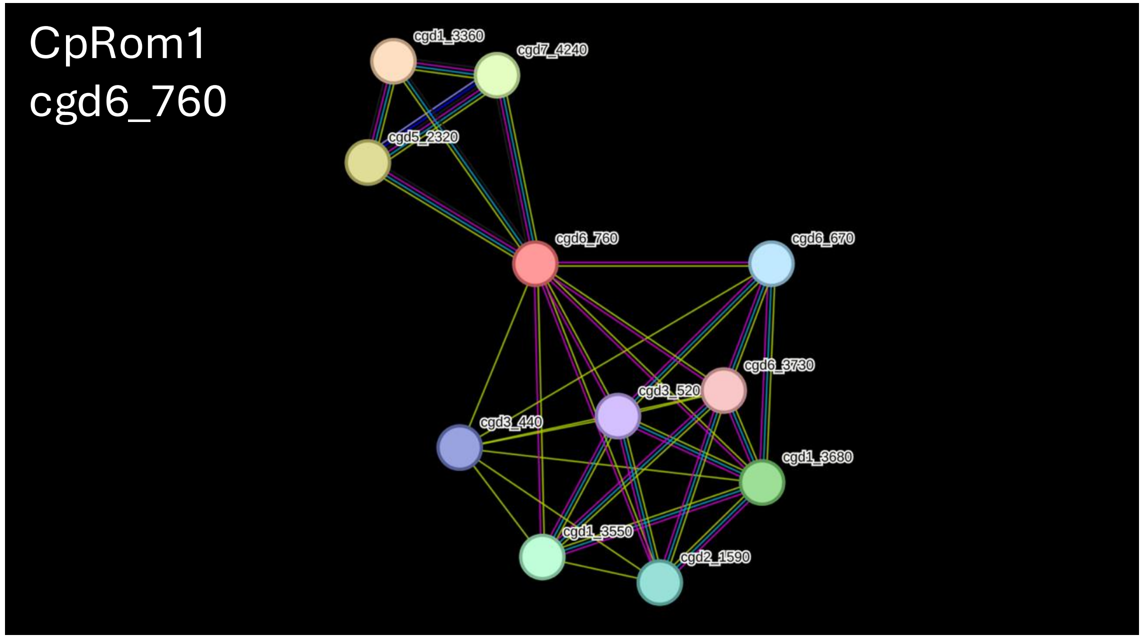


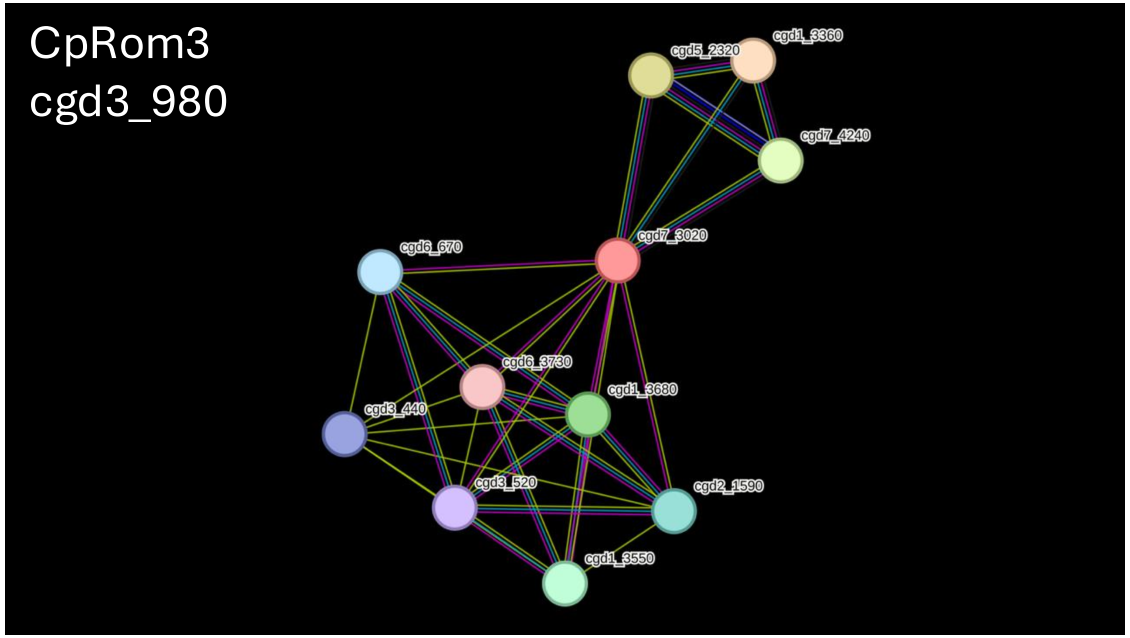


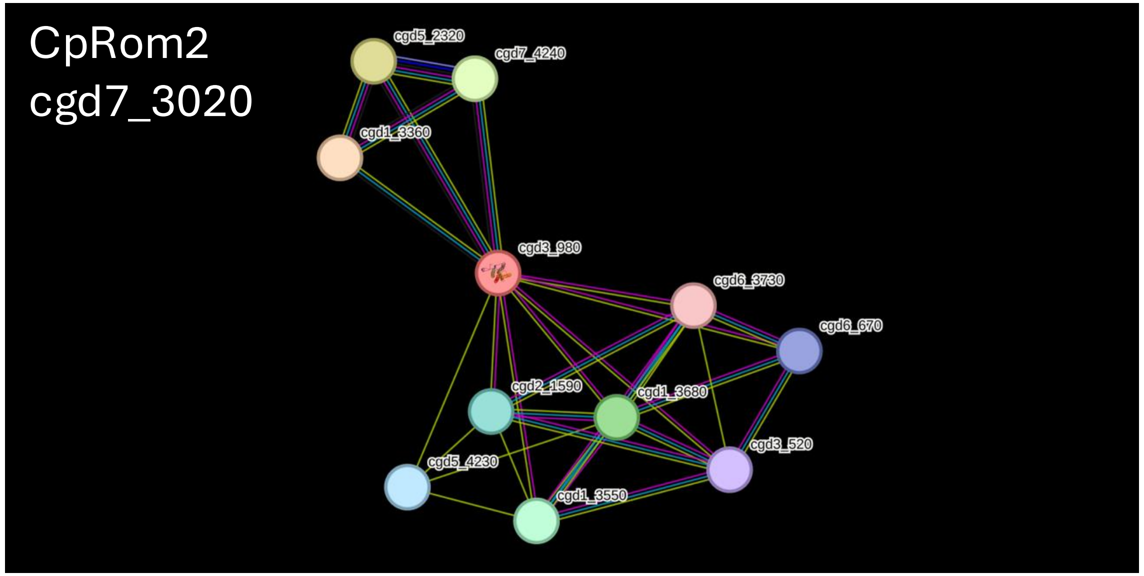


| CryptoDB ID | Interaction | Name |
|---|---|---|
| cgd1_3360 | CpRom1, CpRom2 and CpRom3 | Predicted AFG1 ATpase family AAA ATpase. |
| cgd5_2320 | CpRom1, CpRom2 and CpRom3 | Prohibitin. |
| cgd7_4240 | CpRom1, CpRom2 and CpRom3 | Prohibitin. |
| **cgd1_3680** | CpRom1, CpRom2 and CpRom3 | Extracellular membrane associated protein with 3 EGF domains and a transmembrane domain. |
| cgd1_3550 | CpRom1, CpRom2 and CpRom3 | Mucin-like low complexity glycoprotein with a signal peptide and an apple domain. |
| cgd2_1590 | CpRom1, CpRom2 and CpRom3 | Extracellular protein with signal peptide, 5xEGF and apple domains. |
| cgd6_670 | CpRom1, CpRom2 and CpRom3 | annotation not available |
| cgd3_520 | CpRom1, CpRom2 and CpRom3 | Cysteine-rich extracellular protein with a signal peptide and two apple domains. |
| cgd6_3730 | CpRom1, CpRom2 and CpRom3 | Large extracellular protein with a signal peptide, apple domain and a transmembrane region. |
| cgd3_440 | CpRom1 and CpRom2 | C-type lectin containing protein with a transmembrane domain and mucin-like rich regions |
| cgd5_4230 | CpRom3 | EGF-like domain-containing protein |
