## Supplementary Table 1 for "Exploring the Repertoire of Rhomboid Proteases in *Cryptosporidium parvum* Parasite: Phylogenesis, Structural motifs and Cellular Localization in Sporozoite Cells"

Table 1, of oligonucletides used in this study.

| Gene | Forward primers* | Reverse primers * | Function |
|---|---|---|---|
| CpRom1 | ATGGATATGTCCGATTTTGTTTTC | TCAAGAAAAATCATATCCAAATAC | |
| CpRom1 Nterm | ATGGATATGTCCGATTTTGTTTTC | ATTAACAACTAAACCTCCAAGAGC | |
| CpRom1 Cterm | GGTTGTTCTCCTGAGGATAG | TCAAGAAAAATCATATCCAAATAC | PCR on genomic DNA and RT-PCR on mRNA |
| CpRom2 | ATGTCTGACAGAAAGATTTTTG | TTATCCACATCTTCTAATCCATG | |
| CpRom3 | CACAGACTTTCTGATTTACCTC | CCATACTTGACCCTCCTTAAC | |
| CpRom1 | | TCCC<u>CCCGGG</u>TCATCAAGAAAAATCATATCCAAATA | |
| CpRom1 Nterm | AAC<u>GAGCTC</u>GATATGTCCGATTTTGTTTTCA | TCCC<u>CCCGGG</u>TCAATGTATTGGATTAACCCATTTT | |
| CpRom1 Cterm | AAC<u>GAGCTC</u>TTCTATCCTCCATTATATTGG | TCCC<u>CCCGGG</u>TCATCAAGAAAAATCATATCCAAATA | |
| CpRom2 | AAC<u>GAGCTC</u>TCTGACAGAAAGATTTTTGATAT | | Cloning in expression vector |
| CpRom2 Cterm | AAC<u>GAGCTC</u>TAAGCCATTGTACACTAAGTT | TCCC<u>CCCGGG</u>TTATCCACATCTTCTAATCCAT | |
| CpRom3 | | TCCC<u>CCCGGG</u>TCAAGGATTCATAAGTTTCTCT | |
| CpRom3 Nterm | CGC<u>GGATCC</u>TCAAATATACACAGACTTTCTG | TCCC<u>CCCGGG</u>TTAACTATGTTTCCAAGTGATTCC | |

*Underlined sequences represent restriction sites inserted for cloning.
