## Supplementary file S2 for "Exploring the Repertoire of Rhomboid Proteases in *Cryptosporidium parvum* Parasite: Phylogenesis, Structural motifs and Cellular Localization in Sporozoite Cells"

| | UNIPROT | length | PFAM Peptidase S54 rhomboid | | |
|---|---|---|---|---|---|
| CpRom1 | Q5CXK3 | 990 | 633-769 | 567-587 | 638-657 |
| CmRom1 | B6ABI5 | 892 | 542-678 | 474-494 | 543-561 |
| CpRom2 | F0X3H6 | 464 | 175-309 | 114-131 | 184-206 |
| CmRom2 | B6ABU2 | 469 | 167-300 | 107-123 | 176-198 |
| GnRom1 | A0A023B066 | 350 | 198-333 | 139-158 | 207-228 |
| TgRom5 | Q6GV23 | 841 | 460-602 | 323-343 | 464-484 |
| PfRom4 | Q8I433 | 759 | 457-591 | 332-351 | 466-487 |
| TgRom4 | Q695T8 | 641 | 330-466 | 219-239 | 352-372 |
| GnRom2 | A0A023B0P3 | 448 | 90-229 | 40-58 | 98-122 |
| PfRom3 | A0A5K1K8S0 | 267 | 83-221 | 43-61 | 68-84 |
| TgRom3 | Q6IUY1 | 263 | 79-219 | 37-57 | 86-106 |
| CpRom3 | F0X528 | 282 | 92-230 | 49-71 | 94-116 |
| CmRom3 | B6AAF3 | 273 | 87-226 | 44-66 | 89-111 |
| GnRom3 | A0A023BBG2 | 276 | 94-231 | 50-71 | 96-118 |
| TgRom2 | Q695T9 | 283 | 106-244 | 62-82 | 114-134 |
| PfRom1 | A8IWX2 | 278 | 96-238 | 52-73 | 105-124 |
| TgRom1 | Q695U0 | 293 | 103-247 | 62-82 | 112-132 |
| PfRom10 | C6KT26 | 274 | 98-238 | 55-81 | 139-157 |
| PfRom8 | Q8ILY3 | 738 | 555-698 | 506-529 | 591-611 |
| PfRom7 | A0A5K1K991 | 340 | 170-337 | 188-205 | 211-231 |
| TgRom6 | Q2PP52 | 531 | 349-508 | 277-297 | 307-327 |
| CmRom4 | B6ADY7 | 315 | 92-246 | 50-69 | 150-173 |
| PfRom6 | A0A5K1K8X1 | 569 | 305-478 | 61-70 | 72-81 |
| PfRom9 | Q8I3V7 | 488 | 333-483 | 266-286 | 376-394 |

| | UNIPROT | length | PFAM DER1 Derlin | | |
|---|---|---|---|---|---|
| PfDer1 | Q8IKV3 | 354 | 152-337 | 192-215 | 229-248 |
| TgDer1 | C9WWW6 | 589 | 303-493 | 353-376 | 388-411 |
| PfDer1-2 | Q8IJ82 | 263 | 12-205 | 19-42 | 51-74 |
| PfDerlin-1 | C7SP48 | 214 | 10-202 | 19-42 | 54-77 |
| TgDer1ER2 | C9WWW8 | 212 | 10-200 | 14-37 | 57-80 |

**TM**

| | | | | | |
|---|---|---|---|---|---|
| 669-691 | 697-715 | 727-746 | 904-925 | | |
| 582-600 | 606-623 | 635-654 | 660-681 | 808-831 | |
| 213-233 | 239-258 | 270-288 | 294-314 | 400-419 | |
| 205-225 | 231-254 | 261-280 | 286-307 | 400-419 | |
| 240-257 | 263-282 | 294-311 | 317-341 | | |
| 492-512 | 526-546 | 571-590 | 673-693 | | |
| 521-541 | 553-572 | 578-599 | 681-703 | | |
| 382-402 | 404-424 | 445-465 | 567-587 | | |
| 134-152 | 158-177 | 189-206 | 212-234 | | |
| 96-117 | 129-145 | 151-172 | 184-201 | 207-224 | 236-258 |
| 121-141 | 142-162 | 189-209 | 231-251 | | |
| 128-151 | 157-180 | 189-208 | 214-233 | 245-266 | |
| 123-146 | 152-177 | 184-203 | 209-228 | 240-261 | |
| 130-152 | 158-178 | 190-206 | 218-236 | 248-270 | |
| 149-169 | 179-199 | 205-225 | 227-247 | 260-280 | |
| 136-154 | 166-190 | 196-214 | 221-239 | 254-274 | |
| 148-168 | 174-194 | 217-237 | 262-282 | | |
| 169-189 | 195-214 | 221-238 | 250-272 | | |
| 655-672 | 678-696 | 708-733 | | | |
| 264-286 | 292-311 | 318-337 | | | |
| 367-387 | 407-427 | 440-460 | 484-504 | | |
| 184-207 | 226-245 | | | | |
| 266-275 | 277-286 | 325-336 | 371-389 | 431-448 | 458-476 |
| 403-423 | 427-447 | 461-481 | | | |

**TM**

| | | |
|---|---|---|
| 276-299 | 303-326 | |
| 458-481 | | |
| 94-117 | 120-138 | 144-167 |
| 96-119 | 139-162 | 171-193 |
| 94-117 | 137-160 | 169-189 |
